# Automated model-free analysis of cryo-EM volume ensembles with SIREn

**DOI:** 10.1101/2024.10.08.617123

**Authors:** Laurel F. Kinman, Maria V. Carreira, Barrett M. Powell, Joseph H. Davis

## Abstract

Cryogenic electron microscopy (cryo-EM) has the potential to capture snapshots of proteins in motion and generate hypotheses linking conformational states to biological function. This potential has been increasingly realized by the advent of machine learning models that allow 100s-1,000s of 3D density maps to be generated from a single dataset. How to identify distinct structural states within these volume ensembles and quantify their relative occupancies remain open questions. Here, we present an approach to inferring variable regions directly from a volume ensemble based on the statistical co-occupancy of voxels, as well as a 3D-convolutional neural network that predicts binarization thresholds for volumes in an unbiased and automated manner. We show that these tools recapitulate known heterogeneity in a variety of simulated and real cryo-EM datasets, and highlight how integrating these tools with existing data processing pipelines enables improved particle curation and the construction of quantitative conformational landscapes.

## INTRODUCTION

The proteins and protein complexes that carry out cellular functions are inherently dynamic, occupying high-dimensional conformational landscapes where different free energy minima correspond to different structural and functional states. These landscapes can be altered by environmental signals, drug treatments, and more. Methods to resolve lowly-populated but biologically-informative states within the landscapes, and to quantitatively compare conformational landscapes across different conditions, are therefore instrumental to understanding the function and regulation of this essential cellular machinery. Cryogenic electron microscopy (cryo-EM) is, in principle, a powerful tool to accomplish just this, as standard single-particle analysis (SPA) approaches image individual protein molecules. Assigning each particle image to a structural state affords the potential to build highly detailed, semi-quantitative maps of protein’s structural landscapes, and to ask and answer hypothesis-driven questions about how those landscapes change as a function of experimental condition.

In reality, the power of cryo-EM to analyze these conformational landscapes has been limited. The particle images captured during cryo-EM data collection have extremely low signal-to-noise ratios as a result of the low-dose imaging conditions used to minimize electron beam-induced sample damage (Sigworth 2016). Thus, 10^3^-10^6^ particle images representing potentially highly disparate structural states must be averaged to produce high-resolution three-dimensional reconstructions (Sigworth 2016). Traditional approaches to classifying particles into different structural states have proven widely useful (Scheres *et al*. 2006; Scheres 2012; Zhang *et al*. 2019; Nakane and Scheres 2021) but rely on the assumption that particles can be grouped into *k* discrete classes, where *k* is a relatively small number that users do not know *a priori*, and where different values of *k* or different initial volumes can have substantial impacts on the outputs of these algorithms (Rabuck-Gibbons *et al*. 2022).

Recently, however, several approaches have emerged that leverage the generative power of machine learning (ML) to tackle this problem and begin realizing the single-molecule potential of heterogeneous cryo-EM (Zhong *et al*. 2021; Chen and Ludtke 2021; Punjani and Fleet 2023; Powell and Davis 2024). In particular, approaches like cryoDRGN (Zhong *et al*. 2021), tomoDRGN (Powell and Davis 2024), and e2gmm (Chen and Ludtke 2021) use autoencoder or autoencoder-like architectures to map images of individual particles into a learned latent space that acts as a lower-dimensional representation of the structural heterogeneity in the particle images. Decoding these latent encodings subsequently enables generation of large heterogeneous ensembles of 3D volumes. These approaches have been used to resolve dynamics of radial spoke proteins important for ciliary motility (Gui *et al*. 2021), to identify diverse structural states of ribosomes (Powell and Davis 2024; Sun *et al*. 2023; Kinman *et al*. 2022; Leesch *et al*. 2023), to understand catalysis in a non-ribosomal peptide synthetase (Wang *et al*. 2022), and to quantify subtle structural changes in AAA+ proteases (Ghanbarpour *et al*. 2023a; Ghanbarpour *et al*. 2023b; Ghanbarpour *et al*. 2024). However, questions remain about how to systematically analyze the heterogeneity present in the resulting volume ensembles in order to both identify biologically-interesting modes of variability, and compare this variability quantitatively across ensembles or datasets derived from different experimental conditions.

We have previously shown the power of atomic-model-based approaches to quantitatively characterize the conformational landscape of the assembling bacterial ribosome (Davis *et al*. 2016; Kinman *et al*. 2022; Sun *et al*. 2023); however, such approaches are limited in several key respects. First, they require an atomic model to be fit within each map in an ensemble, so datasets where there is no existing atomic model, or where the volumes are so heterogeneous that an atomic model could not be well fit into all volumes, cannot be analyzed using this approach. Furthermore, model-based approaches assume that a feature is either present in its modeled location, or absent; as such, they are challenged by conformational heterogeneity, where they often report a “partial” native occupancy. In such instances, we have previously shown that complementary approaches such as principal component analysis of the volume ensembles are valuable in characterizing large-scale conformational changes (Sun *et al*. 2023). Lastly, model-based approaches are fundamentally hypothesis-driven, as they require users to supply a mask indicating where within the three-dimensional volume they expect to detect heterogeneity. While this approach can be powerful, particularly when paired with orthogonal biochemical or genetic evidence that drives the structural hypothesis, it prevents the user from detecting heterogeneity in unexpected regions of the structure.

To address these limitations, and to aid users in detecting heterogeneity within volume ensembles without requiring a model, we have developed SIREn (<u>S</u>ubunit Inference from <u>R</u>eal-space <u>En</u>sembles). Our SIREn tool is not to be confused with methods for 3D reconstruction based on sinusoidal representation networks (SIRENs) (Sitzmann *et al*. 2020) recently reported by the Carazo and Sorzano groups (Herreros *et al*. 2024). SIREn leverages the co-variance of voxel occupancies across volumes within the ensemble to detect structural subunits without the aid of an atomic model. SIREn thus identifies putative structural subunits by measuring how voxels are co-occupied across the ensemble, and by extracting groups of voxels that are highly co-occupied. We show that SIREn successfully extracts blocks corresponding to compositional and conformational heterogeneity we encoded in simulated datasets (*i*.*e*., ground truth), as well as in several well-studied, highly heterogeneous real datasets from both SPA pipelines and *in situ* tomographic data. Furthermore, we showcase the development of a 3D-convolutional neural network (3D-CNN) to predict contour levels across volume ensembles, and we demonstrate the integration of SIREn with our previously developed MAVEn tool (Sun *et al*. 2023) to both curate particle stacks for high-resolution refinement, and quantify the abundance of conformational states of interest.

## RESULTS

### SIREn algorithm design

Recent advances in ML-enabled heterogeneous reconstruction methods permit hundreds-to-thousands of density maps to be generated from a single cryo-EM or cryogenic electron tomography (cryo-ET) dataset. We hypothesized that we could leverage the statistical power of these large volume ensembles to directly infer regions within the volume that vary across the ensemble, which we refer to as structural blocks. Specifically, we asserted that voxels belonging to the same structural block should be positively and negatively co-occupied (both occupied or both unoccupied within a given volume, respectively) more often than expected by chance, whereas voxels belonging to distinct structural blocks were expected to be co-occupied at a rate predicted by their overall rates of occupancy across the ensemble. Diverse structural states of a protein complex could then be defined by isolating sets of particles with similar patterns of occupancy across these fundamental building blocks (**Figure 1A**).

**Figure 1.**
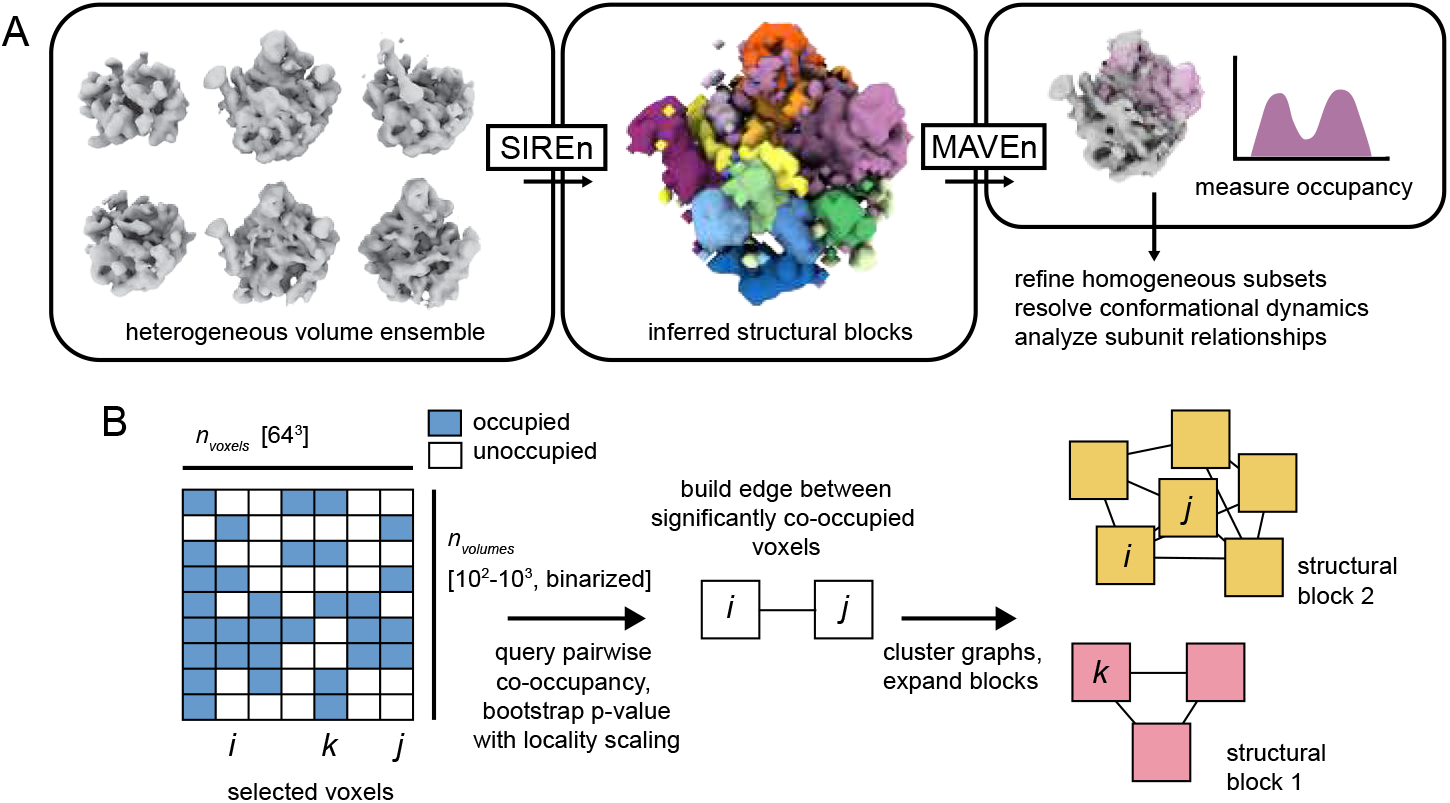
Inferring structural heterogeneity with SIREn. **(A)** Flowchart depicting application of SIREn to infer variable regions (structural blocks) directly from a heterogeneous volume ensemble, and downstream application of SIREn blocks to query occupancy of features of interest using MAVEn. **(B)** Schematic overview of the SIREn algorithm. Illustrative heatmap depicting binarized volumes as rows within the depicted array, with each voxel determined to be occupied or unoccupied within a given volume based on a binarization threshold provided for each map. Voxels are queried for co-occupancy, and a graph is constructed with edges connecting voxels determined to be significantly co-occupied. Clustering and expansion of the graph (see Methods) produces the final SIREn-identified blocks (pink and yellow, right).

To detect these blocks, we employed a two-step approach. In step one, we queried pairs of voxels for frequency of positive and negative co-occupancy across an ensemble of binarized volumes, and used a bootstrapping method to determine whether the measured co-occupancy was likely to be significant (**Figure 1B**). We then constructed a graph with nodes corresponding to voxels, and edges connecting voxels related by the bootstrapped significance measured above, and subsequently clustered the graph using a label propagation approach (Cordasco and Gargano 2010) to produce “seed blocks”. In the second step, the seed blocks were expanded using a similar bootstrapping method: if a given voxel exhibited significant co-occupancy to a minimum fraction of the seed voxels in a block, we added it to the block. In this manner, we queried each voxel against each seed block to produce the final structural blocks (see Methods).

Given that we do not expect the underlying volume data to be independent and identically distributed, we emphasize that we employed this bootstrapping method not as a rigorous statistical test but rather as a heuristic for estimating the strength of the evidence that the occupancies of two voxels were related. In practice, we additionally modified these calculations to include a “locality scaling factor” for positive co-occupancy that depended on the physical distance between the two voxels (**Supplemental Figure 1A**, see Methods). The inclusion of this scaling factor reflected the intuition that physically neighboring voxels are more likely to be constituents of the same block, and thus less stringent statistical evidence should be demanded to cluster those voxels. In contrast, distal voxels, which could represent tightly coupled allosteric structural changes, should require substantially stronger statistical evidence to be assigned to the same block. Notably, our two-step approach to infer structural blocks permitted voxels to be added to more than one structural block, an important feature given that we expected conformational motions to result in several distinct but overlapping blocks.

#### A 3D-convolutional neural network enables unbiased prediction of contour levels across volume ensembles

SIREn takes as input a large (100s-1,000s) ensemble of volumes aligned to a single reference frame. These volumes must be binarized at a provided threshold value, which is equivalent to setting a contour level. Defining appropriate binarization thresholds for such a large number of volumes is a challenging task: statistical approaches, such as binarizing at the 99th percentile of the volume data as is standard in ChimeraX (Meng et al. 2023), often produce poor binarization thresholds for any given volume, and manual assignment of thresholds represents both a throughput challenge and a potential source of bias. To address these challenges, we noted that the Electron Microscopy Data Bank (EMDB) (Turner et al. 2024) represents a large, well-curated training dataset containing diverse structural density maps alongside their annotated binarization thresholds. We hypothesized that a 3D-convolutional neural network (3D-CNN) trained on these volumes and annotations could predict appropriate thresholds for unseen maps, and we therefore retrieved, curated, and pre-processed ∼ 4,000 such maps to train such a 3D-CNN (see Methods).

We next constructed a 3D-CNN consisting of five convolutional layers upstream of a multi-layer perceptron (MLP), adapted from a previously implemented architecture (Lee *et al*. 2022). This model was designed to output a single binarization threshold for each input map (**Figure 2A**, see Methods). The trained network successfully returned thresholds highly correlated with the expert-curated labels deposited in the EMDB in both the training set and the withheld validation and test sets. Notably, it outperformed the often-used 99^th^ percentile standard using metrics of Pearson correlation coefficient and proximity to a one-to-one relationship between predicted and annotated thresholds (**Figure 2B, Supplemental Figure 2A-B**). Individual exemplar maps visualized at the threshold predicted by the 3D-CNN reinforced that the predicted labels are consistent with the ground truth and outperform statistical approaches (**Figure 2B**). Interestingly, in the withheld test and validation sets, we found that the 3D-CNN predicted thresholds effectively distinguished between protein and micellar density for maps of membrane proteins (**Supplemental Figure 2C**), consistent with the network predicting thresholds that emphasize the shapes observed in protein, DNA, and RNA macromolecular complexes common in the EMDB.

**Figure 2.**
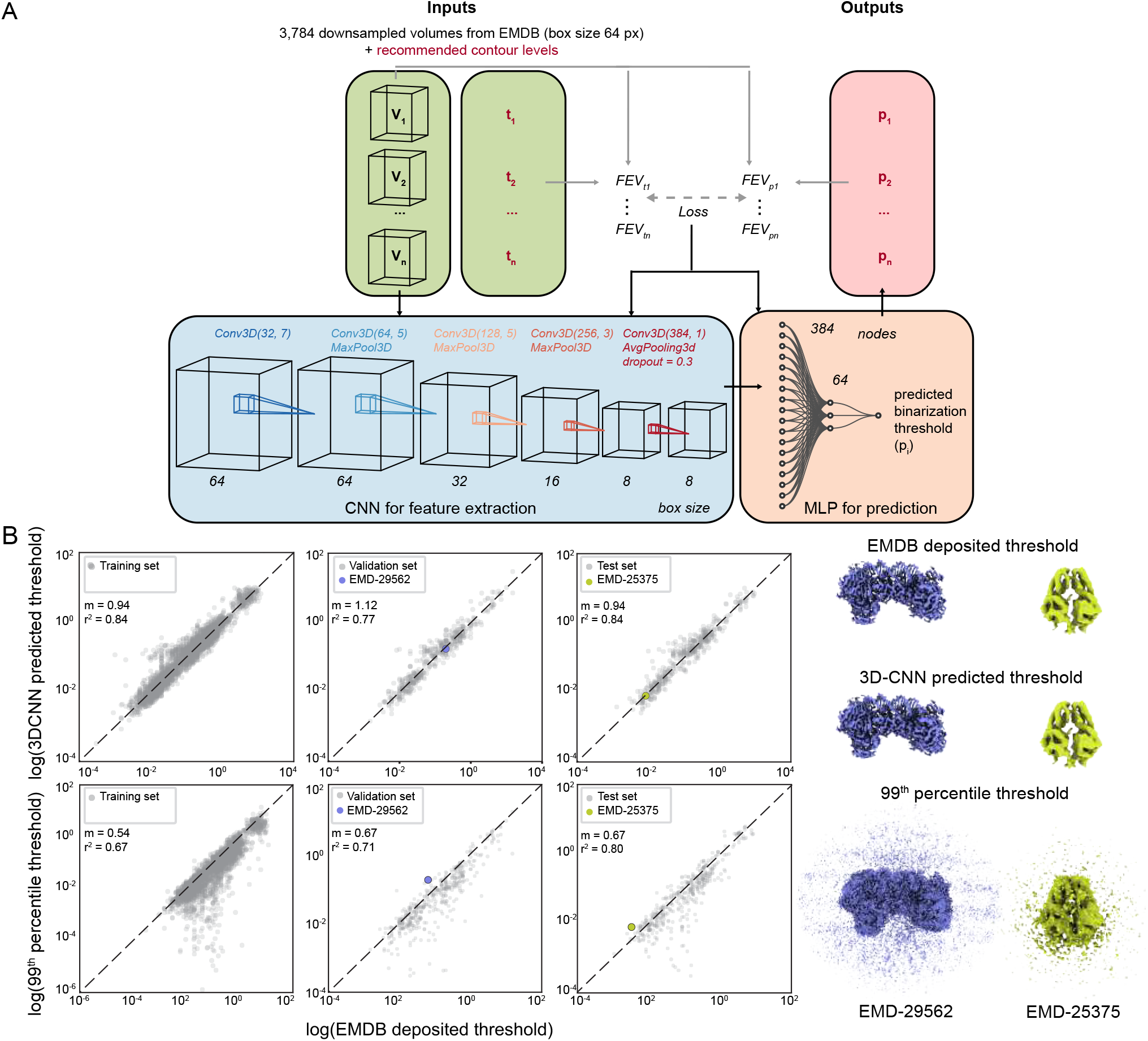
3D-convolutional neural network (3D-CNN) enables accurate prediction of binarization thresholds at scale. **(A)** Depiction of the 3D-CNN inputs (green box), model architecture (blue box – 3D-CNN; peach box -MLP), and outputs (red box). Gray arrows indicate information used to calculate the training loss from the fraction enclosed volume (FEV) at input (ti) or predicted (pi) binarization thresholds (see Methods). **(B)** Scatter plots comparing EMDB ground truth labels and 3D-CNN-predicted thresholds (top), or thresholds predicted by the 99th percentile of the data (bottom) for training (left), validation (middle), and test (right) sets. Slopes (m) and Pearson correlation coefficients (r2) are noted over each plot; the identity line is shown as a dashed black line on each plot. Exemplar maps EMD-29562 (Wang *et al*. 2023; blue) and EMD-25375 (Fry *et al*. 2022; green) (right) are shown at EMDB-deposited (top), 3D-CNN predicted (middle), and 99th percentile (bottom) binarization thresholds.

### SIREn robustly detects compositional heterogeneity in simulated SPA datasets

To assess whether SIREn could detect structural heterogeneity within a volume ensemble, and to probe the limits of this detection, we next constructed a series of simulated cryo-EM single particle analysis datasets consisting of the bacterial large ribosomal subunit (50S) with some or all of the protein uL2 removed (see Methods). Particle images of the 50S subunit with uL2 intact (uL2^Δ0^) were then titrated at various frequencies into particle stacks containing different uL2 deletions (*i*.*e*., uL2^Δ0.25^, uL2^Δ0.50^, uL2^Δ0.75^, uL2^Δ1.0^) to yield a total of twelve simulated particle stacks with varying heterogeneity. We also generated five homogeneous particle stacks, corresponding to each of the five constructs employed in building the heterogeneous particle stacks (**Figure 3A**, see Methods).

**Figure 3.**
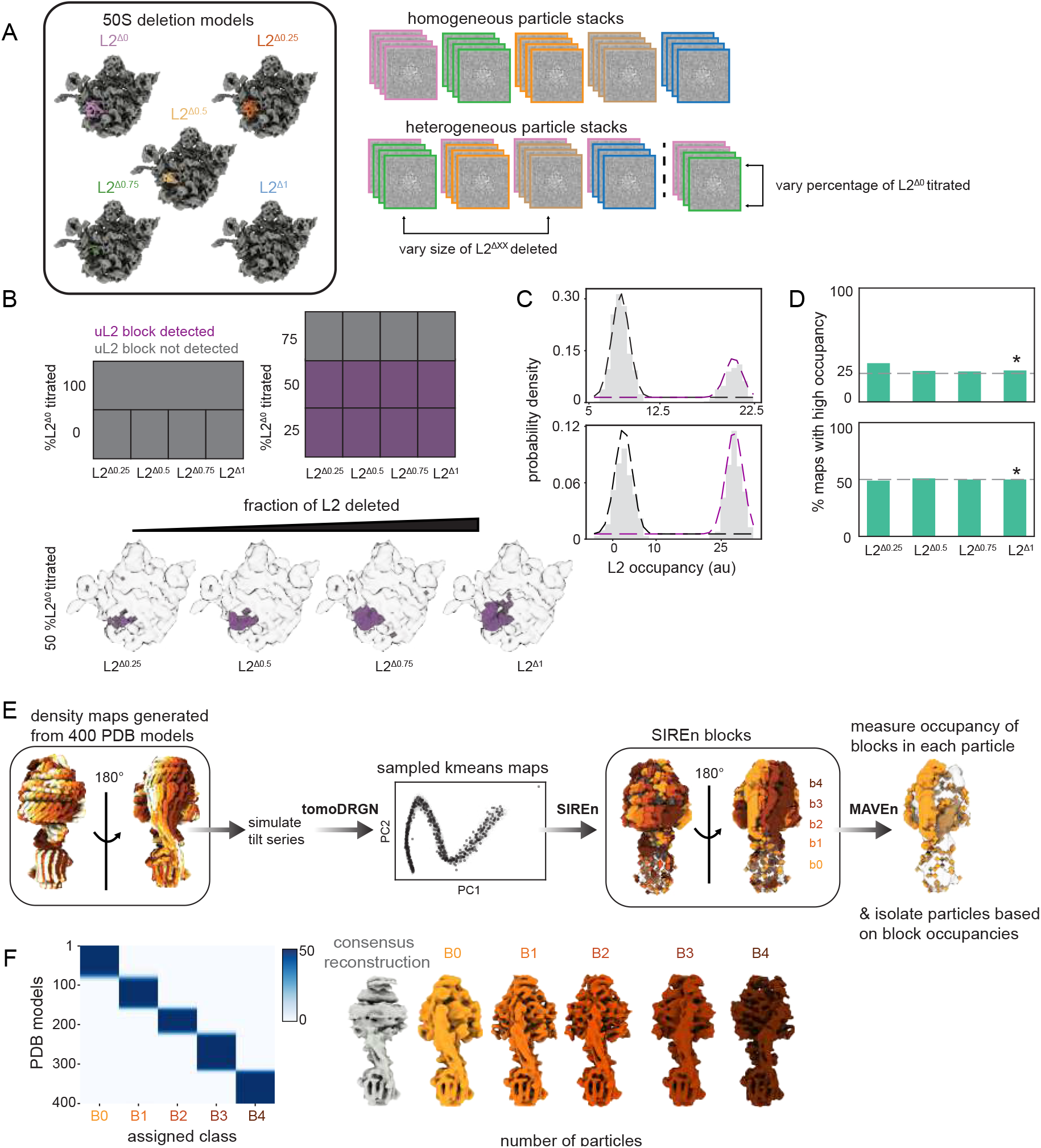
SIREn recapitulates ground truth compositional and conformational heterogeneity in simulated datasets. **(A)** Atomic models (left) of the *E. coli* large ribosomal subunit (50S) with full (L2Δ1) or partial (L2^Δ0^, L2^Δ0.25^, L2^Δ0.50^, L2^Δ0.75^) deletion of uL2, with uL2 highlighted in various colors. A schematic overview (right) of the generated homogeneous and heterogeneous simulated particle stacks, titrating both the proportion of uL2 deleted and the fraction of particles with uL2 intact, is shown (right). Particle images are outlined with colors matching the 50S deletion model panel. **(B)** Heatmaps (top) depicting whether a uL2 block was detected by SIREn in each dataset. The detected uL2 blocks (bottom) are shown in purple for the four datasets containing 50% uL2-intact (L2^Δ0^) particles, and 50% particles bearing successively larger deletions of uL2 (L2^Δ0.25^ – L2^Δ1^). Blocks are overlaid on an exemplar density map from the ensemble (transparent surface). **(C)** Representative distributions of per-volume uL2 block occupancies in datasets with 25% (top) or 50% (bottom) uL2 intact, with the remainder of the particles having no uL2 (uL2^Δ1^). Dashed lines indicate the fit two-component Gaussian mixture model (GMM) used to distinguish low-occupancy particles (black) from high-occupancy particles (purple, see Methods). **(D)** Fraction of uL2-intact particles in each dataset where a uL2 block was detected, as determined by the GMM fitting approach (see Methods). Asterisks indicate the datasets highlighted in C. The ground truth frequency is shown as a horizontal dashed line in the barplots. **(E)** Workflow to analyze a simulated cryo-ET dataset of yeast ATP synthase, with atomic models traversing the simulated conformational change colored yellow-to-red (left), and locations of 500 maps sampled from latent space, projected using principal component analysis. Detected SIREn blocks are shown in shades of orange and red (middle). A representative map from the ensemble (translucent surface) with overlaid SIREn-detected block (orange) is shown to illustrate occupancy-querying approach with MAVEn. **(F)** Confusion matrix comparing the inferred class assignments and ground truth rotational state for each particle in the simulated particle stack (left). Reconstructed volumes corresponding to each class are shown (right). Volumes are colored to match blocks in E.

We trained a series of cryoDRGN models on the resulting datasets and generated a volume ensemble consisting of 500 maps at box size 64 for each dataset. These volume ensembles were used as the inputs for SIREn. As expected, SIREn did not identify a uL2 block in the homogeneous volume ensembles, where particles were derived from a single atomic model. SIREn successfully identified a full or partial uL2 block in all heterogeneous ensembles consisting of 25% or 50% uL2-intact particles (**Figure 3A**), but failed to do so when analyzing volume ensembles resulting from particle stacks with 75% uL2-intact particles. We attribute the asymmetry of this detection – that a block was detected at 25% uL2-intact particles but not 75% uL2-intact particles – to the fact that the SIREn algorithm requires more stringent evidence for positive co-occupancy than negative co-occupancy, making it challenging to detect blocks for highly-occupied features.

Satisfyingly, in those cases where SIREn did identify a block, the size of the block scaled with the proportion of uL2 deleted (**Figure 3A, Supplemental Figure 3A**), suggesting that SIREn successfully identified the correct spatial extent of the heterogeneity in the volumes. Notably, SIREn was able to detect a block even when only 25% of the uL2 protein was deleted. This perturbation corresponds to ∼8 kDa (73 amino acids), relative to the total 50S ribosomal mass of ∼1.3 MDa, suggesting that, despite downsampling to box size 64, with the resolution Nyquist-limited at 6.9 Å, SIREn can detect relatively subtle structural changes.

We hypothesized that we could use the structural blocks inferred by SIREn to estimate the frequency of uL2-intact particles in each volume ensemble. To do so, we used the detected structural blocks directly as input masks in MAVEn, a software we have previously developed for quantifying the occupancy of structural subunits across large volume ensembles (Kinman *et al*. 2022; Sun *et al*. 2023). We fit the resulting occupancy distributions of the uL2 structural block in each volume ensemble to a two-component Gaussian mixture model, and assigned each volume to either a low-or high-uL2 occupancy class (**Figure 3C, Supplemental Figure 3B**, see Methods). We compared the number of particles assigned to the high uL2 occupancy class to the known frequency of uL2-intact particles in each simulated dataset, finding close correspondence between these values **(Figure 3D**), which indicated that our pipeline can accurately quantify the number of particles in a given conformational state.

### SIREn identifies conformational heterogeneity in a simulated tomographic dataset

To determine whether SIREn could similarly detect conformational heterogeneity within a volume ensemble generated using cryo-ET, we used an existing simulated tomographic dataset (Powell and Davis 2024) of yeast ATP synthase in diverse rotational states (Guo and Rubinstein 2022). Briefly, Powell and Davis simulated 20,000 tilt-series image sets from density maps generated using 400 different atomic models representing a continuous conformational trajectory, and used these tilt series to train a tomoDRGN model (Powell and Davis 2024). We sampled 500 density maps from the latent space of the trained tomoDRGN model, and supplied these volumes as the input ensemble to SIREn. SIREn detected five blocks that highlighted the coupled rotations of the stalk and the nucleotide binding site (**Figure 3E**). To isolate particles representing different rotational states, we coupled our 3D-CNN-predicted binarization thresholds with MAVEn (Sun *et al*. 2023) to generate a volume for every particle in the input dataset, and queried that volume ‘on-the-fly’ for occupancy of each of the SIREn-annotated blocks (**Supplemental Figure 4A**). Notably, incorporating our 3D-CNN-predicted binarization thresholds allowed us to control for per-volume scaling effects, and ensured that block occupancies scaled between 0 and 1 (see Methods). Based on the resulting occupancy measurements, we were able to assign each particle to one of the five rotational states identified by SIREn (**Figure 3F, Supplemental Figure 4B**, see Methods). Finally, we compared our inferred particle class assignments to the known (*i*.*e*., ground truth) rotational state of each particle, finding that our combined SIREn-MAVEn pipeline accurately identified the rotational state of each particle at this 5-class level of discretization (**Figure 3F**).

**Figure 4:**
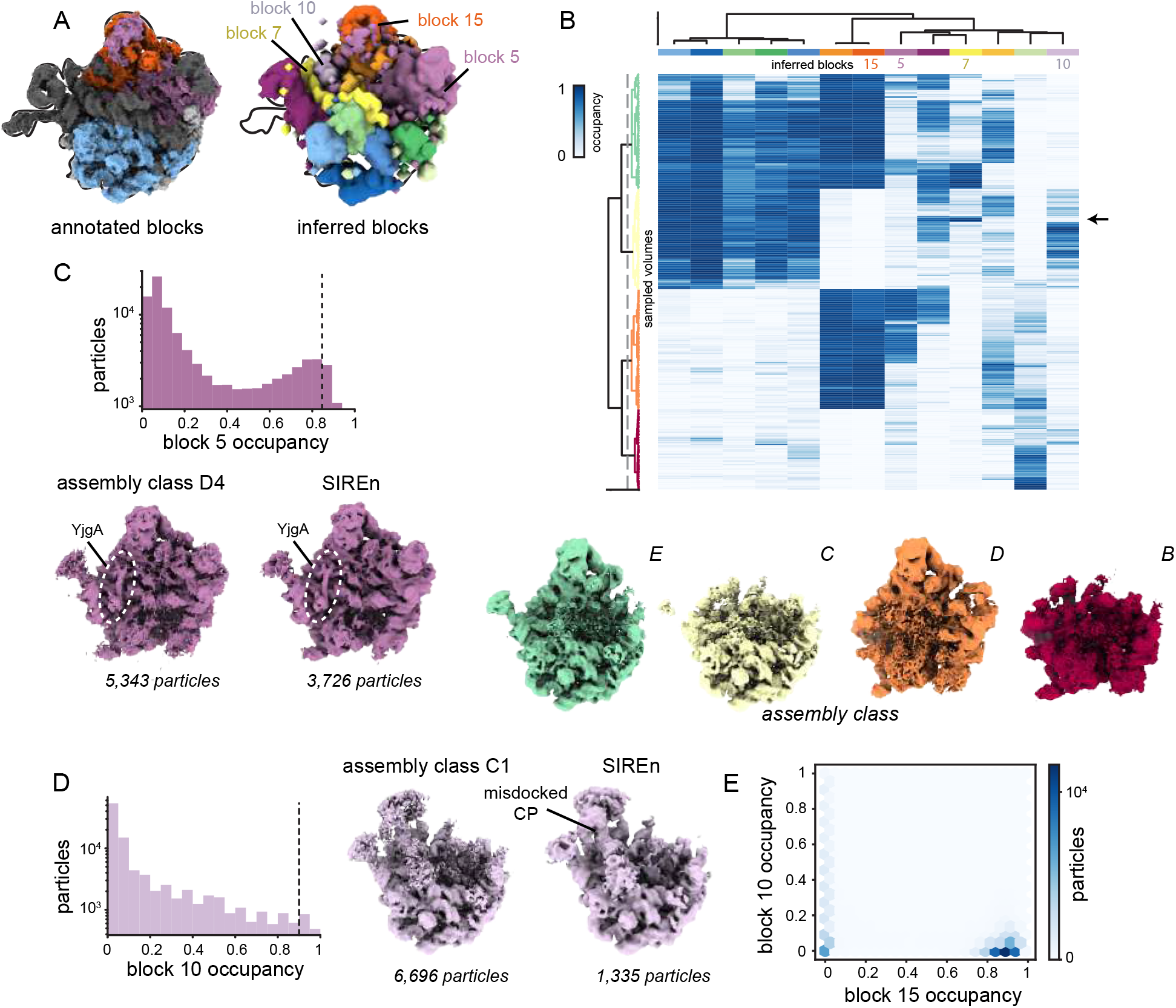
SIREn identifies variable features in real cryo-EM datasets. **(A)** Blocks inferred by SIREn (right) from the EMPIAR-10076 dataset of assembling bacterial ribosomes (Davis *et al*. 2016), compared to blocks annotated (left) by a model-based approach in a previous analysis (Kinman *et al*. 2022) of this dataset. **(B)** Heatmap depicting occupancy of inferred blocks (columns of heatmap) in each of the 500 maps used as inputs to SIREn (rows of heatmap). Columns are colored to match (A). Rows were clustered to generate four major classes, from which density maps were generated (see Methods)and annotated following Davis *et al*. 2016. The arrow marks volumes with high occupancy for block 7, which corresponds to helix 68, but low occupancy for block 15, which corresponds to the central protuberance. **(C)** Results from querying the full particle stack for occupancy of block 5 (see Methods), depicted as a histogram. Dashed line indicates the threshold used to select particles for homogeneous refinement. Density map resulting from a refinement performed with SIREn-curated particles compared to the D4 assembly class map (right) reported by Davis *et al*. 2016. Dashed white line surrounds YjgA. **(D)** Results from querying the full particle stack for occupancy of block 10, depicted as a histogram (left) with dashed line indicating threshold used to select particles for homogeneous refinement. Density map resulting from refinement performed on SIREn-curated particles compared to the published C1 assembly class map (right) reported by Davis *et al*. 2016. **(E)** A density map depicting the joint distribution of block 10 (misdocked CP) and block 15 (properly docked CP) occupancies obtained by the ‘on-the-fly’ querying of the volume ensemble (see Methods).

To assess if such inferred particle classes could improve the consensus refinement and resolve different structural states across the range of motion sampled by ATP synthase in this dataset, we performed reconstructions in RELION (Zivanov *et al*. 2018) on each of the inferred particle classes. As expected, these reconstructions were higher quality than the consensus reconstruction and resolved distinct rotational states of the complex (**Figure 3F**).

### SIREn identifies known assembly states of the bacterial large ribosomal subunit

Given that SIREn successfully identified known heterogeneity in simulated datasets, we were interested in benchmarking its performance on a well-annotated real cryo-EM dataset – specifically a dataset of the assembling 50S ribosome, isolated from a bL17-depletion strain (EMPIAR-10076) (Davis *et al*. 2016). This dataset has previously been characterized by both extensive 3D classification (Davis *et al*. 2016; Rabuck-Gibbons *et al*. 2022), model-based analysis of cryoDRGN-generated volume ensembles (Kinman *et al*. 2022), and other reconstruction methods aimed at visualizing structural heterogeneity (Chen and Ludtke 2021; Punjani and Fleet 2021; Gilles and Singer 2024). Each of these methods revealed four primary structural states and a series of additional sub-states.

To determine whether SIREn would identify the known structural blocks in this dataset, we employed the same cryoDRGN model previously trained on this dataset (Zhong *et al*. 2021; Kinman *et al*. 2022), and sampled 500 volumes at box size 64 from the resulting latent space via *k*-means clustering. The inferred SIREn blocks closely corresponded to the structural blocks identified by the MAVEn atomic-model-based approach, with blocks corresponding to several features known to have variable occupancy across the ensemble, including the central protuberance, base, uL1 stalk, bL12 stalk, and H68 (**Figure 4A**). We calculated the normalized occupancy of each block in the input volume ensemble, with each volume binarized at the threshold predicted by the 3D-CNN, and clustered the results, permitting us to identify groups of structurally-related volumes (**Figure 4B**). We identified four major classes of particles from this clustering, which closely match the B-E classes identified by previous analyses (Davis *et al*. 2016); volumes generated at the centroid position in latent space of each of the four volume classes confirmed that these particles represented the B-E classes (**Figure 4B**). SIREn additionally highlighted interesting intra-class heterogeneity within each of these four primary classes, including a small number of volumes with occupancy for H68 but not the central protuberance (block 15, **Figures 4A-B**). This “C4 class” was not observed in the original hierarchical classification analysis, and was first identified in a previous cryoDRGN analysis that relied on the atomic model for interpretation (Zhong *et al*. 2021).

Interestingly, we observed that the SIREn block corresponding to the bL12 stalk also included density proximal to the binding site for the ribosome biogenesis factor YjgA (block 5, **Figure 4A**), which was unexpectedly first identified via 3D classification in the original analysis of this dataset (Davis *et al*. 2016). To determine whether particles with high occupancy of this block represented the YjgA-bound state, we performed the same ‘on-the-fly’ querying approach described above, wherein we generated a volume at box size 64 for every particle in the input stack, binarized these volumes according to their 3D-CNN-predicted thresholds, and measured the occupancy of the resulting volumes. We then selected ∼3,700 particles with high occupancy of this block and used these particles to perform homogeneous reconstruction in cryoSPARC. The resulting reconstruction bore strong density for YjgA (**Figure 4C**).

In addition to the YjgA binding site block, we observed another block near H68 that was not well explained by the atomic model (block 10, **Figure 4A**). By similarly querying the occupancy of every particle for block 10, and selecting ∼1,300 particles with high block 10 occupancy, we were able to clearly resolve the central protuberance in a misdocked conformation (Leidig *et al*. 2014). Notably, the density for the misdocked central protuberance in the reconstruction from our SIREn-identified particles is better defined than the corresponding density in the originally reported C1 class volume, despite our reconstruction using substantially fewer particles (**Figure 4D**). Lastly, our approach allowed us to directly compare the relationship between occupancy of the properly-docked central protuberance (block 15) and the misdocked central protuberance (block 10); as expected, occupancy of these two features was mutually exclusive (**Figure 4D**).

### Uncovering intra- and inter-molecular heterogeneity of the bacterial ribosome *in situ* with SIREn

To understand how SIREn performed on real cryo-ET data, we applied tomoDRGN and SIREn to analyze *Mycoplasma pneumoniae* ribosomes imaged *in situ* (EMPIAR-10499) (Tegunov *et al*. 2021). Briefly, a tomoDRGN model was previously trained on ribosomes extracted from tomograms of chloramphenicol treated *M. pneumoniae* cells, and careful inspection of the tomoDRGN-generated volume ensemble revealed extensive heterogeneity within the 70S ribosomes, including highly occupied P- and A-tRNA sites and minor states with the E-site tRNA occupied (Powell and Davis 2024). To compare the structural blocks inferred by SIREn to these previous manual annotations, we provided SIREn with 500 density maps sampled by *k*-means clustering of the latent space from the tomoDRGN model. This automated SIREn analysis corroborated conformational and compositional heterogeneity in the ribosome previously identified by expert-guided inspection (**Figure 5A**). Specifically, blocks highlighting heterogeneity in the uL1 (block 19) and bL12 (block 6) stalks were detected, as well as a block near the tRNA A-site (block 20). To more deeply interrogate these sites, we again turned to our ‘on-the-fly’ querying approach (see Methods) to select particles with high occupancy of each of these blocks (**Figure 5A**, right). We performed reconstructions using the high-occupancy particles for each block in RELION and compared the resulting maps with either maps reconstructed using low-occupancy particles or a consensus reconstruction using the entire particle stack. The resulting maps clearly resolved differential occupancy of each of these features (**Figure 5A**).

**Figure 5:**
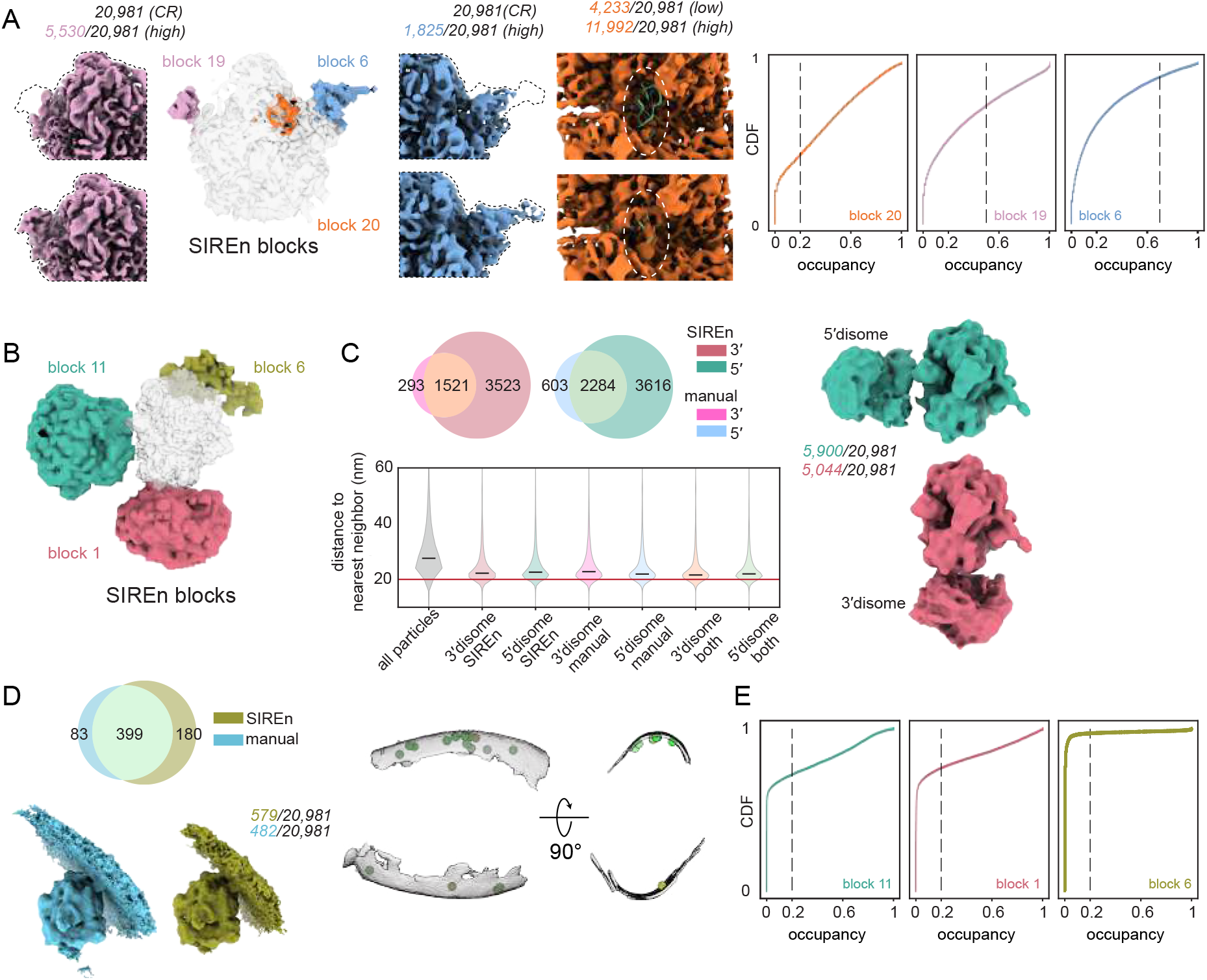
SIREn uncovers heterogeneous features of cellular ribosomes in cryo-ET datasets. **(A)** Blocks detected by SIREn in an ensemble of 500 ribosomes resolved by cryo-ET from *M. pneumoniae* cells (EMPIAR-10499). Insets compare the consensus reconstruction using all particles (CR) and particle subsets with either high or low occupancy of the noted block. Maps are colored to match the block used to filter the particle stack, and the location of variability is highlighted with a dashed line. The number of particles used for each reconstruction is listed in italics, and PDB model 7phb (Xue *et al*. 2022) is docked in the block 20 reconstruction (green). Particle occupancies determined by ‘on-the-fly’ querying (see Methods) are shown as cumulative density functions (CDF), and thresholds used to identify high-occupancy particles are noted with dashed lines (right). **(B)** Blocks identified by SIREn from a volume ensemble of ribosomes extracted at larger real-space box size. **(C)** Venn diagrams depicting total number of particles labeled as 5’ or 3’ disomes by SIREn (Supplemental Figure 4A, see Methods) and by published manual annotations (top). Reconstructions performed with SIREn-annotated 5’ and 3’ disome particles (right); number of particles used in each reconstruction listed. Distribution of nearest-neighbor distances in sets of particles annotated by different methods (SIREn, manual, or both; bottom). **(D)** Venn diagram comparing manual and SIREn annotations for membrane-bound ribosomes (top). Reconstructions performed using particles from manual (blue) and SIREn (yellow) annotations are shown (bottom). Numbers of particles used for each reconstruction listed. Membrane-annotated particles in a representative tomogram are shown (right). Segmented membrane density is shown in gray (see Methods), particles are colored as at left. **(E)** Cumulative density function plots of the per-particle occupancy of disome and membrane blocks (see Methods). Thresholds used for particle selection are shown as dashed lines.

The unique potential of performing cryo-ET on intact cells or milled lamellae is that it confers the power to not only resolve structures *in situ* but also investigate the spatial and cellular context of these structures. As a result, we were interested in determining whether SIREn could be applied to understand intermolecular heterogeneity in the cellular milieu surrounding ribosomal particles. To test this, we sampled 500 maps from the latent space of a tomoDRGN model trained on particles extracted with a larger real-space box (∼74 nm, as opposed to ∼36 nm for the particles used to train the previous model) (Powell and Davis 2024). After running SIREn on this volume ensemble, we observed two blocks that we hypothesized represented 5’ and 3’ disomes (blocks 11 and 1, Figure 5B), as well as a block that likely corresponded to membrane density (block 6, Figure 5B). We again defined an empirical threshold to isolate particles with high occupancy of each block and used RELION to perform a reconstruction with these particles (Zivanov et al. 2018) (Figure 5E, see Methods). We identified extensive overlap between our SIREn-based automated annotations and those generated manually (Powell and Davis 2024), with SIREn notably capturing a larger number of both disomes and membrane-associated ribosomes. We verified that the additional particles we annotated are likely to represent true disomes in two ways: first, we calculated the distance of each annotated particle to its nearest neighbor, and compared the distribution of nearest-neighbor distances in SIREn- and manually-annotated particles, finding both sets had smaller average nearest-neighbor distances than unannotated particles (**Figure 5C**). Secondly, we performed reconstructions using the particles identified by SIREn as 5’ and 3’ disomes, and found that doing so revealed low-resolution ribosomal density adjacent to the well-resolved central monosome (**Figure 5C**). Similarly, reconstructions using SIREn-identified membrane-associated particles resolved membrane density, and we found that the particles SIREn annotated as membrane-associated mapped to the membrane in source tomograms (**Figure 5D**).

## DISCUSSION

This work contributes two tools to aid in the automated, model-free analysis of highly heterogeneous cryo-EM volume ensembles: a 3D-CNN that predicts an appropriate binarization threshold given an input density map, and a bootstrapping and graph clustering approach for inferring regions of heterogeneity directly from a volume ensemble. These tools fill a key niche in modern cryo-EM processing workflows as they help guide interpretation of the large number of volumes produced by modern generative reconstruction algorithms (Zhong *et al*. 2021; Chen and Ludtke 2021; Punjani and Fleet 2021; Gilles and Singer 2024), or from deep classification with traditional tools (Rabuck-Gibbons *et al*. 2022). Specifically, we show how these tools can help to resolve minor states and quantify the frequency of specific conformational or compositional features across both SPA and cryo-ET datasets.

### Automated prediction of density map binarization thresholds with a 3D-CNN

Choosing accurate binarization thresholds (equivalently, contour levels or isosurface levels) for 3D density maps is both essential, and challenging to do at scale and in an unbiased manner. Choice of binarization threshold impacts which features are visible in a volume, and thus substantially impacts analysis of maps generated by EM data processing pipelines. Historically, binarization thresholds have been set to the 99^th^ percentile of the data or manually defined, however, the need for more automated approaches to predicting binarization thresholds has been noted (Beckers *et al*. 2019; Joseph *et al*. 2020). A previous method to solve this problem by Pfab and Si, which used a minimization approach based on fraction of voxels above the threshold and the surface area-to-volume ratio, highlighted the utility of an unbiased method for threshold prediction (Pfab and Si 2019).

Here, we address this challenge by leveraging publicly available data on the EMDB (Turner *et al*. 2024) to train a 3D-convolutional neural network (3D-CNN) to predict contour levels for density maps. To our knowledge, ours is the first machine-learning-based approach to predicting binarization thresholds for cryo-EM density maps, although CNNs have proven useful in other cryo-EM data processing tasks (Giri *et al*. 2023; de Teresa-Trueba *et al*. 2023; Mu *et al*. 2021). Although we apply the 3D-CNN specifically to predict thresholds for SIREn, we anticipate that this tool will be broadly useful in visualizing and analyzing 3D density maps generated by cryo-EM or cryo-ET. For example, there is a need for increased standardization of map thresholds, particularly as presented in publications, and all maps presented here are displayed at the threshold predicted by the 3D-CNN. Moreover, any approaches to volume ensemble analysis that entail measuring and comparing the occupancy of various subunits, particularly across independently collected datasets, will likely benefit from the appropriate application of binarization thresholds. Indeed, such comparisons otherwise require careful intra-dataset normalizations (Sun *et al*. 2023). To facilitate broad adoption of the 3D-CNN, we have developed a publicly available script (https://github.com/mariacarreira/calc_level_ChimeraX) that allows users to rapidly predict volume thresholds directly in ChimeraX (Meng *et al*. 2023).

### Inferring structural blocks with SIREn

In addition to an unbiased method for predicting contour levels for cryo-EM density maps, we also present SIREn, a tool for directly inferring regions of heterogeneity within a large structural ensemble. There are several approaches one could take to this task, all of which rely on examining how voxel occupancies co-vary across the dataset. Indeed, a previously-published approach (Sheng *et al*. 2023) involves performing dimensionality reduction with PCA and UMAP (McInnes *et al*. 2018), followed by clustering with HDBSCAN (Campello *et al*. 2015). One could similarly classify voxels into blocks via hierarchical clustering using simple correlation metrics, or other clustering-based approaches to partition voxels into non-overlapping classes. In contrast, SIREn allows voxels to be assigned to more than one class, which enables the detection of overlapping blocks. Our approach furthermore allows voxels that may have relatively weak correlations, but statistically significant co-occupancies, to be classified together. SIREn’s design additionally allows us to impose prior knowledge about the real-space relationship between voxels within a subunit by implementing a locality scaling factor that effectively prioritizes identifying relationships between physically proximal voxels. Notably, we only apply this scaling factor to positive co-occupancy, and generally employ weaker statistical constraints in identifying significant negative co-occupancy so as not to overly penalize voxels that may participate in multiple structural blocks.

In practice, we find that SIREn performs well in identifying known heterogeneity within datasets: SIREn is able to identify features like a misdocked central protuberance and a bound assembly factor in a well-annotated real dataset of assembling bacterial ribosomes, which we could not have identified using atomic-model-based approaches like those we have previously described (Sun *et al*. 2023).

Nonetheless, there remain limitations to the performance of SIREn, the first being that input volumes must typically be substantially downsampled. Although we are able to detect changes as small as ∼8 kDa in the simulated uL2 deletion datasets, interesting smaller scale heterogeneity may be lost due to this downsampling. This limitation follows from the combinatorial scaling in the number of calculations SIREn performs in querying pairs of voxels for co-occupancy; even relatively minor increases in the number of non-background voxels (caused by either a larger box size, or a larger solvent mask) substantially impact SIREn running time (**Supplemental Tables 1-2**). SIREn running time is also impacted by the size of the volume ensemble, as ensemble size informs the dimensions of the precalculated look-up table used to determine significance thresholds. In our tests, ensemble sizes in the range of 500-1,000 volumes have performed well. Finally, the size of blocks that can be detected is also limited by the random subsampling we perform in the initial step of block detection. We perform this random subsampling to limit the multiple hypothesis testing burden and to decrease run time. We find that subsampling by a larger factor than the default value of 2 generally decreases sensitivity, though users may wish to adjust this hyperparameter based on available computational resources.

We also emphasize that SIREn is, first and foremost, a hypothesis generation tool, and we recommend validating observations through traditional high-resolution refinement to identify which of the detected blocks are likely to represent biologically-meaningful heterogeneity. We present several semi-automated pipelines for performing such analysis, including methods for querying occupancy of each identified block for every particle in a dataset, and performing traditional homogeneous refinements using subsets of particles identified in this manner. We showcase the power of this approach in our analysis of an assembling bacterial large ribosomal subunit dataset, where we are able to resolve YjgA and the misdocked central protuberance. In particular, for both YjgA and the misdocked CP, our homogeneous refinements have more interpretable density for these features than the original maps produced by 3D classification, despite using substantially fewer particles (**Figure 4C-D**), suggesting that our SIREn-MAVEn approach is able to curate a more homogeneous subset of particles than 3D classification produces.

### Benchmarking tools for heterogeneity analysis

To benchmark the performance of SIREn, we used a combination of simulated datasets and well-annotated real datasets. Using simulated datasets, with their precise ground-truth annotations, we were able to probe the limits of SIREn’s detection capabilities. We found that identifying features that were highly occupied across a dataset, such as those seen in the datasets with uL2 titrated at 75% frequency (**Figure 3A**), was challenging, but that SIREn was nonetheless able to detect variable features as small as ∼8 kDa in the context of the ∼1.3 MDa 50S ribosome. Similarly, we found that SIREn was able to resolve distinct blocks corresponding to the continuous conformational trajectory sampled by the yeast ATP synthase dataset (**Figure 3C**). Although our approach necessarily discretizes a continuous trajectory, we were able to use our ground-truth labels to reveal that SIREn correctly inferred labels assigning each particle to a distinct rotational state (**Figure 3D**).

How well simulated datasets recapitulate the features of real datasets for the purposes of benchmarking heterogeneity analysis tools, however, remains an open question (Noble 2024). In particular, real datasets contain features we do not see in simulated datasets that contribute to error and noise, including ice artifacts, radiation damage, incorrect orientation assignments from the 3D refinement procedure, and more (Baxter *et al*. 2009). Moreover, it is not clear that the common noise models applied to generate simulated data capture the effects of noise in real datasets (Baxter *et al*. 2009; Himes and Grigorieff 2021). To address these limitations, we also benchmark our tools using several real, well-annotated datasets (**Figures 4-5**), noting that in these real datasets, we do not have access to ground-truth labels for the conformational state of each particle, which limits our ability to precisely examine algorithmic performance. We therefore highlight that there is a potential broad utility for the development of better tools to accurately benchmark the performance of heterogeneous reconstruction algorithms and downstream analysis tools.

### Resolving heterogeneity *in situ* with SIREn

We developed SIREn with highly heterogeneous SPA datasets in mind, but we found that it could be readily applied to cryo-ET datasets. This is perhaps unsurprising given that the SIREn algorithm is largely agnostic to upstream processing and relies simply on having a large ensemble of 3D density maps aligned to a common reference frame. The observation that SIREn performs well for cryo-ET data as well as SPA data is exciting, as it affords the possibility of quantitatively exploring conformational landscapes *in situ*. For example, here we showed SIREn could detect differential occupancy of the A-site tRNA and bL12 and uL1 stalks in ribosomes imaged directly within intact *M. pneumoniae* cells (Tegunov *et al*. 2021). The potential for broader application of these approaches is still limited by our ability to perform robust particle picking and refinement on non-ribosomal particles *in situ*, but there is promising progress towards that end with optimized lamella production (Khavnekar *et al*. 2022), new particle picking approaches (Rice *et al*. 2023), and large cryo-ET datasets in which sub-nanometer resolution structures of non-ribosomal particles have begun to be solved (Khavnekar *et al*. 2023). In the long term, we envision applying SIREn and related approaches to map heterogeneous structures back to their native contexts to better understand how these structures interact with the physical landscape of the cell.

### Automated tools for the quantitative analysis of conformational landscapes

As structural biology moves increasingly towards characterizing proteins as they exist in cells – occupying highly dynamic conformational landscapes, engaging in diverse transient interactions, and existing within distinct subcellular spatial contexts – we envision a broad utility for approaches like SIREn that enable systematic and quantitative analysis of highly heterogeneous structural ensembles. Beyond simply identifying where structural heterogeneity occurs within a 3D volume, we show here that SIREn-MAVEn can be used to quantitatively query the occupancy of inferred blocks across a particle stack. Having done so, we can not only curate subsets of particles that bear a feature of interest, but also examine patterns of block occupancy to uncover structural interdependencies. For example, we use these quantitative measurements to demonstrate that occupancy of the misdocked CP and properly-docked CP are, as expected, mutually exclusive (**Figure 4D**). Applying similar approaches to datasets where an experimental condition has been varied – for example, treatment with a drug or the addition of a binding partner – should allow one to interrogate how structural ensembles respond to specific perturbations.

## METHODS

### SIREn algorithm design

The SIREn algorithm is built on the intuition that two voxels that belong to the same structural block should be both occupied and both unoccupied in a given volume more often than predicted by their rates of occupancy across the ensemble. To formalize these intuitions, we consider two voxels *v*_*i*_ and *v*_*j*_, occupied at frequencies *f*_*i*_ and *f*_*j*_ in an ensemble of binarized volumes. If the occupancy of *v*_*i*_ and *v*_*j*_ is independent, then we expect the probability that *v*_*i*_ and *v*_*j*_ are both occupied in a volume **V** to be given by

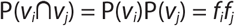

and similarly that

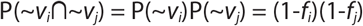

describes the probability of neither voxel being occupied.

To estimate the expected spread of values around the center under the assumption that occupancy of *v*_*i*_ and *v*_*j*_ are not related, we use a bootstrapping approach. For every unique pair of frequencies *f*_*i*_ and *f*_*j*_ found in the input ensemble, we computationally resample with replacement to generate a new ensemble with randomly-distributed occupancies of the voxels *v*_*i*_ and *v*_*j*_. We perform this resampling operation 1,000 times, and use the results to define a cut-off threshold *t*_*ij*_ at which co-occupancy of the voxels occurs less frequently than a defined p-value. To control for multiple hypothesis testing, we employ a Bonferroni correction on this p-value. If we observe that P(*v*_*i*_∩*v*_*j*_)>*t*_*ij*_ and P(∼*v*_*i*_∩∼*v*_*j*_)>*t*_*∼i∼j*_, we determine that the occupancies of *v*_*i*_ and *v*_*j*_ are not independent, and *v*_*i*_ and *v*_*j*_ are candidates to belong to the same structural block. To reduce the multiple hypothesis testing burden and increase computational efficiency, these calculations are performed only on voxels occupied in at least 1% of the volume ensemble; all voxels occupied less frequently than in 1% of volumes are excluded as solvent background. Moreover, we randomly subsample the voxels in this initial querying step, performing these bootstrap calculations on every pair of voxels from a randomly-selected half of the remaining non-background voxels. Finally, when determining whether a given pair of voxels are significantly co-occupied, we modify the positive co-occupancy threshold by a factor *c*_*ij*_≥1 that we dub the ‘locality scaling factor’ and which is dependent on the Euclidean distance between *v*_*i*_ and *v*_*j*_ (**Supplemental Figure 1**), so that we consider *v*_*i*_ and *v*_*j*_ to be related if P(*v*_*i*_∩*v*_*j*_)>*c*_*ij*_\**t*_*ij*_.

After using bootstrap approaches to estimate the strength of the evidence for each pair of voxels being related, we then construct a graph where the nodes are the voxels, and where an edge is built between each pair of voxels *v*_*i*_ and *v*_*j*_ that are determined to be significantly related. We use a label propagation graph clustering method (Cordasco and Gargano 2010) to generate clusters of voxels that serve as seed blocks. Having generated the seed blocks, we then query each voxel *v*_*i*_ against each seed voxel *s*_*k,j*_ in a given seed block *S*_*k*_ using the bootstrapping approach described above. If *v*_*i*_ is significantly related to a minimum fraction of the seed voxels (empirically, we have found 25% to be a reasonable threshold) in *S*_*k*_, we add *v*_*i*_ to *S*_*k*_. Blocks are saved as binary volumes in MRC format, suitable for directly visualizing in ChimeraX (Meng *et al*. 2023), and using as input masks to MAVEn (Sun *et al*. 2023).

### Neural network design and architecture

Input maps used for training the 3D-CNN were downloaded from the Electron Microscopy Data Bank (EMDB) (Turner *et al*. 2024). A total of 5,627 maps were downloaded from the EMDB, along with their corresponding annotated contour levels, based on structure determination method (single-particle cryo-EM) and resolution (<10 Å). These entries were filtered to select maps with box sizes less than 916 pixels and fractional enclosed volumes (see below) ranging from 0.001 to 0.02, resulting in 4,730 entries. Test and validation sets (10%) were selected at random, resulting in a final training set of 3,784 maps that were Fourier cropped to a box size of 64 pixels. Following Ranno and Si (see eqs. 1-3), the maps and their accompanying labels were normalized based on map statistics (Ranno and Si 2022). The scatter plots shown in Figure 2 exclude a small number of maps that had negative binarization thresholds predicted by the 99^th^ percentile approach (60, 5, and 7 maps from the training, validation, and test sets, respectively).

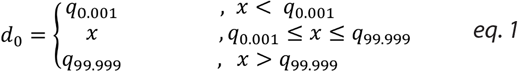

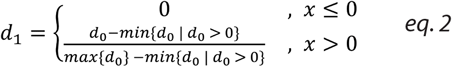

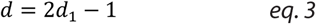

where *q*_*i*_ denotes the i^th^ percentile of the data matrix *x*. The preprocessed (downsampled and normalized) maps were input to a 3D-CNN coupled to a multilayer perceptron (MLP). The architecture of the convolutional layers is adapted from Lee and colleagues (Lee *et al*. 2022), with one batch normalization layer added after each of the first four convolutional layers. The output of the convolutional layers was subjected to a 3D average pooling layer and dropout of 0.3 was applied. The final binarization threshold was predicted by an MLP with the following architecture: a first layer with 384 input and 64 output nodes, a ReLU layer, and a final layer with 64 input nodes and 1 output node. The model was trained for 10 epochs with batch size 32, Adam optimizer, learning rate of 5×10^−6^, and weight decay of 0. Loss during training was calculated by comparing the fraction of the volume enclosed at the predicted threshold – *i*.*e*. what fraction of the voxels in a volume have a value equal to or greater than the predicted threshold – to the fraction of the volume enclosed at the annotated threshold. As such a ‘counting’ operation is not differentiable, we adapted the loss by implementing a soft threshold using a sigmoid function, and calculating the mean squared error between the outputs of this sigmoidal function for the predicted threshold and the annotated threshold (eq. 4). The loss is thus given by:

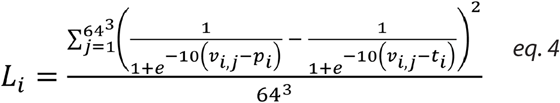

where *L*_*i*_ is the loss calculated by this method for a given map *i*; *p*_*i*_ and *t*_*i*_ are the predicted and annotated labels, respectively, for the map *i*; and *v*_*ij*_ is the intensity of voxel *j* in map *i*.

### Simulating uL2 ribosomal SPA datasets

Full or partial uL2 deletion models were generated by deleting 271, 213, 136, 73, or 0 amino acids from the C-terminus of protein uL2 in the atomic model of the *E. coli* ribosomal large subunit (PDB: 4ybb) (Noeske *et al*. 2015), resulting in the uL2^Δ1^, uL2^Δ0.75^, uL2^Δ0.5^, uL2^Δ0.25^, and uL2^Δ0^ models, respectively. These atomic models were converted to 3D density maps using the `pdb2mrc` command implemented in EMAN2 (Tang *et al*. 2007) at a box size of 440 pixels and pixel size of 1 Å/px. We generated 100,000 noiseless 2D projections from each model, randomly sampling projection angles from SO(3), and randomly sampling a uniform translation distribution with an upper bound of 22 pixels using the `project3d` script as implemented in cryoSRPNT (https://github.com/bpowell122/cryoSRPNT) and described previously (Powell and Davis 2024). Structural and shot noise were added to the projections using the `acn` script in cryoSRPNT, adapted to calculate the standard deviation of a clean particle stack on 2,000 particles at a time, rather than the full particle stack. We subsequently corrupted the resulting particle images via the contrast transfer function (CTF) using the same script. Defocus values for CTF corruption were randomly sampled from a series of Gaussian distributions with means ranging between 10,000 and 25,500 Å, stepped by 5,000 Å, each with a standard deviation of 1,000 Å. The other parameters used in CTF corruption were as follows: spherical aberration of 2.7 nm, microscope voltage of 300 kV, amplitude contrast ratio of 0.1, and phase shift of 0°.

To generate the five homogeneous particle stacks, we used the `write_starfile` script to write a RELION 3.0 .star file for each dataset. To generate the heterogeneous particle stacks, we implemented a `shuffle_particles` script (available with SIREn) to randomly select and shuffle entries from the intact (uL2^Δ0^) and full or partial deletion (uL2^Δ1^, uL2^Δ0.25^, uL2^Δ0.5^, uL2^Δ.25^) uL2 particle stacks and generate a .star file with particles mixed at the desired frequencies. A total of 17 (5 homogeneous and 12 heterogeneous) datasets were generated. For the purpose of training cryoDRGN models, CTF and pose parameters were parsed from the .star files using cryoDRGN’s `parse_ctf_star` and `parse_pose_star` commands, respectively, and particle images were downsampled to a box size of 128 pixels using cryoDRGN’s `downsample` tool. The model architectures used were as follows: encoder and decoder dimensions of 3 layers with 1024 nodes each. Models were trained for 50 epochs with batch size 8. The input volume ensemble for SIREn was generated by performing *k*-means clustering on the cryoDRGN latent encodings with *k* = 500, and generating a volume at each *k*-means cluster center. cryoDRGN v1.1.2 was used for all trained models.

### SIREn analysis of simulated uL2 ribosomal SPA datasets

The input volume ensemble was downsampled and normalized using the SIREn `preprocess` command, which implements the downsampling and normalization described above. The 3D-CNN predicted per-map binarization thresholds for all 500 maps using the SIREn `eval_model` command and the trained model weights. The `sketch_communities` command was run with default p-value thresholds for positive and negative co-occupancy (0.01 and 0.05, respectively) and with random subsampling by a factor of 2. The `expand_communities` command was likewise run with default p-value thresholds for positive and negative co-occupancy (0.01 and 0.05, respectively). In the datasets where an uL2 block was identified, the resulting block was used as mask for MAVEn’s `calc_occupancy` script (https://github.com/lkinman/MAVEn) to query the occupancy of that block in each of the 500 cryoDRGN-generated maps. Occupancy values were normalized by uL2 block size. We fit a two-component Gaussian mixture model (GMM) with tied covariance matrices to the resulting uL2 block occupancy distribution in each dataset, and assigned each volume to a low-or high-uL2 occupancy class based on the fit GMM to quantify the frequency of uL2-intact particles in each dataset.

### SIREn analysis of a simulated cryo-ET dataset

The simulated yeast ATP synthase dataset was generated from PDB models 7tk6, 7tk7, 7tk8, 7tk9, 7tka, 7tkb, 7tkc, and 7tkd (Guo and Rubinstein 2022), and a tomoDRGN model was trained on the resulting data, as described previously (Powell and Davis 2024). Maps (500) were generated at a box size of 104 pixels from the trained model’s latent space using *k*-means clustering, and input to SIREn. Maps were downsampled and normalized, and binarization thresholds were predicted, using the `preprocess` and `eval_model` commands implemented in SIREn as described above. The `sketch_communities` and `expand_communities` commands were run with default values to generate putative structural blocks. The `calc_occupancy_otf` script implemented in MAVEn was adapted to incorporate binarization predictions for each generated map using the 3D-CNN we developed, and to be compatible with tomoDRGN-generated maps. We used this adapted script to query the occupancy of the five blocks representing rotational states of the yeast ATP synthase in each of the 20,000 particles in the dataset. Occupancies were normalized by input block size to scale between 0 and 1 for each block. Each particle was then assigned to an inferred particle class corresponding to the block for which that particle had maximal occupancy. The `filter_star` script from tomoDRGN was used to filter the image series .star file based on indices of particles assigned to each class. The resulting filtered .star files were used to perform reconstructions (without pose refinement) in RELION 3.1. The maps were postprocessed in RELION and low-pass filtered to 10 Å.

### SIREn analysis of EMPIAR-10076

A cryoDRGN (v0.3.2) model was trained on the EMPIAR-10076 dataset, as described previously (Zhong *et al*. 2021; Kinman *et al*. 2022), and 500 maps were sampled from the latent space using *k*-means clustering. Maps were generated at box size 64, and normalized using SIREn’s `preprocess` command. Binarization thresholds were predicted using the `eval_model` command and trained model weights, and structural blocks were predicted using the `sketch_communites` and `expand_ communities` commands with default values. The occupancy of the resulting blocks was measured in each of the 500 input maps, using the predicted binarization thresholds and normalizing occupancies by the size of each block. These occupancy measurements were hierarchically clustered to generate the heatmap shown in Figure 4D. Each of the 500 input maps was assigned to a volume class by implementing a threshold on the dendrogram resulting from the hierarchical clustering; all particles within a *k*-means class were assigned the same volume class as their corresponding *k*-means cluster center volume. The median position in latent space was then calculated for each of the four volume classes, and a centroid volume generated at the nearest on-data point in the latent space. The structural blocks generated by SIREn were also used as input masks in a version of the `calc_occupancy_ otf` script implemented in MAVEn (Sun *et al*. 2023) that was adapted to incorporate SIREn-predicted binarization thresholds for each volume, producing measurements of the occupancy of each block in each of the particles in the dataset (96,478 in total). Particles with high occupancy of block 5 (bL12 stalk and YjgA) and 10 (misdocked central protuberance) were isolated and used to perform homogeneous refinements in cryoSPARC (Punjani *et al*. 2017).

### SIREn analysis of EMPIAR-10499

TomoDRGN models were trained on the EMPIAR-10499 (Tegunov *et al*. 2021) dataset of chloramphenicol-treated *M. pneumoniae* cells as described previously, with one model trained on particles extracted with a real box size of ∼36 nm (the intramolecular heterogeneity model) and another model trained on particles with a real box size of ∼74 nm (the intermolecular heterogeneity model) (Powell and Davis 2024). We performed *k*-means clustering on the latent space of each model, with *k* = 500, and generated maps at box size 64 at each *k*-means cluster center. The resulting maps were low-pass filtered to 8 and 12Å, respectively. These maps were then normalized using SIREn’s `preprocess` command, and binarization thresholds were predicted using the `eval_model` command and trained model weights. The `sketch_communities` command was run with default values after filtering maps with predicted binarization thresholds more than two standard deviations from the mean, thereby excluding low-quality maps deriving from low-quality particles. This capability is directly implemented in the `sketch_communities` and `expand_communites` commands using the `--filter` flag. For both the intramolecular and intermolecular models, the adapted `calc_occupancy_otf` script described above for use with tomoDRGN and SIREn-predicted binarization thresholds was used to generate a map for each particle in the dataset (20,981 particles in total) and to measure occupancy of each block in the resulting maps. Occupancies were normalized by block size to scale between 0 and 1, and subsets of particles were selected as having high or low occupancy of each block using empirically defined cutoffs. In the intramolecular heterogeneity dataset, particles with occupancy of 0 for the relevant blocks were designated as ‘low occupancy’ particles, while thresholds of 0.5, 0.7, and 0.2 were used to select ‘high occupancy’ particles for blocks 19, 6, and 20, respectively. In the intermolecular heterogeneity dataset, all particles with occupancy of membrane or disome blocks greater than 0.2 were designated as ‘high occupancy’ for the relevant block (**Figures 5A**,**E**). The relevant particle subsets were saved as .pkl files and used to filter the appropriate image series .star file using the tomoDRGN `filter_star` script. Filtered .star files were used as inputs for RELION reconstructions (without pose refinement). Maps were post-processed and low-pass filtered to 10 Å (20 Å for the disome maps). SIREn-annotated particles were compared to the manual annotations described by Powell and Davis. Relevant particles were mapped onto tomograms using tomoDRGN’s `subtomo2chimerax` command. Membrane density in the tomograms was segmented using MemBrain-Seg (Lamm *et al*. 2024). Segmented membrane density was postprocessed using a custom jupyter notebook (Powell *et al*. 2024) and ChimeraX (Meng *et al*. 2023).

### Runtime Experiments

For runtime experiments, SIREn was run on a cluster equipped with an NVIDIA GeForce RTX 3090 GPU (24 GB) and two Intel Xeon Gold 6242R CPUs. The resulting runtimes are reported in **Supplemental Tables 1-2**. In cases where the input volumes were not at a box size of 64 pixels, as required for siren `eval_model`, maps were downsampled by siren `preprocess` prior to normalization. Selected real datasets were run using the ‘`--filter`’ argument in the `sketch_communities` and `expand_communities` commands, which filters volumes with outlier predicted binarization thresholds. For the dataset where box size was less than 64 pixels, a single, manually selected binarization threshold was supplied to the `sketch_communities` and `expand_communities` commands.

## CODE AVAILABILITY

The SIREn software is available at: www.github.com/lkinman/SIREn.

Script to automate threshold predictions in ChimeraX using the trained 3D-CNN are available at: www.github.com/mariacarreira/calc_level_ChimeraX.

Scripts to simulate particle stacks are available at: https://github.com/bpowell122/cryoSRPNT.

## AUTHOR CONTRIBUTIONS

Conceptualization: LFK, MVC, JHD.

Investigation: LFK, MVC.

Software: LFK, MVC.

Data Curation: LFK, MVC, BMP.

Writing–original draft: LFK, MVC.

Writing–review and editing: LFK, MVC, BMP, JHD.

Visualization: LFK, MVC, JHD.

Supervision: LFK, JHD.

Project administration and funding acquisition: JHD.

## ACKNOWLEDGEMENTS

We thank Mira May, Alireza Ghanbarpour, Jose-Maria Carazo, and Carlos Oscar Sorzano for their feedback. Work in the Davis lab is supported by funding from the Smith Foundation Odyssey Award, the MIT Jameel Clinic for Machine Learning in Health, NIH grant R01-GM144542, and NSF CAREER grant 2046778.

## SUPPLEMENTARY FIGURES

**Supplemental Figure 1:**
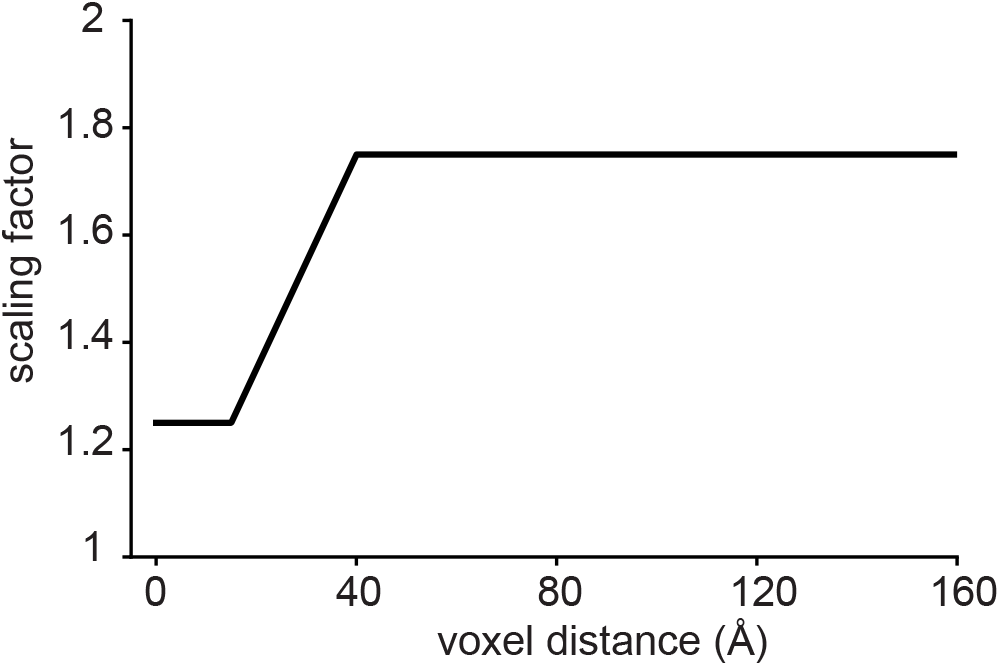
Distance-dependent locality scaling factor. The scaling factor used to modify threshold for positive co-occupancy is shown as a function of voxel distance. Note that the scaling factor remains fixed at 1.75 for all voxel distances greater than 40 Å.

**Supplemental Figure 2:**
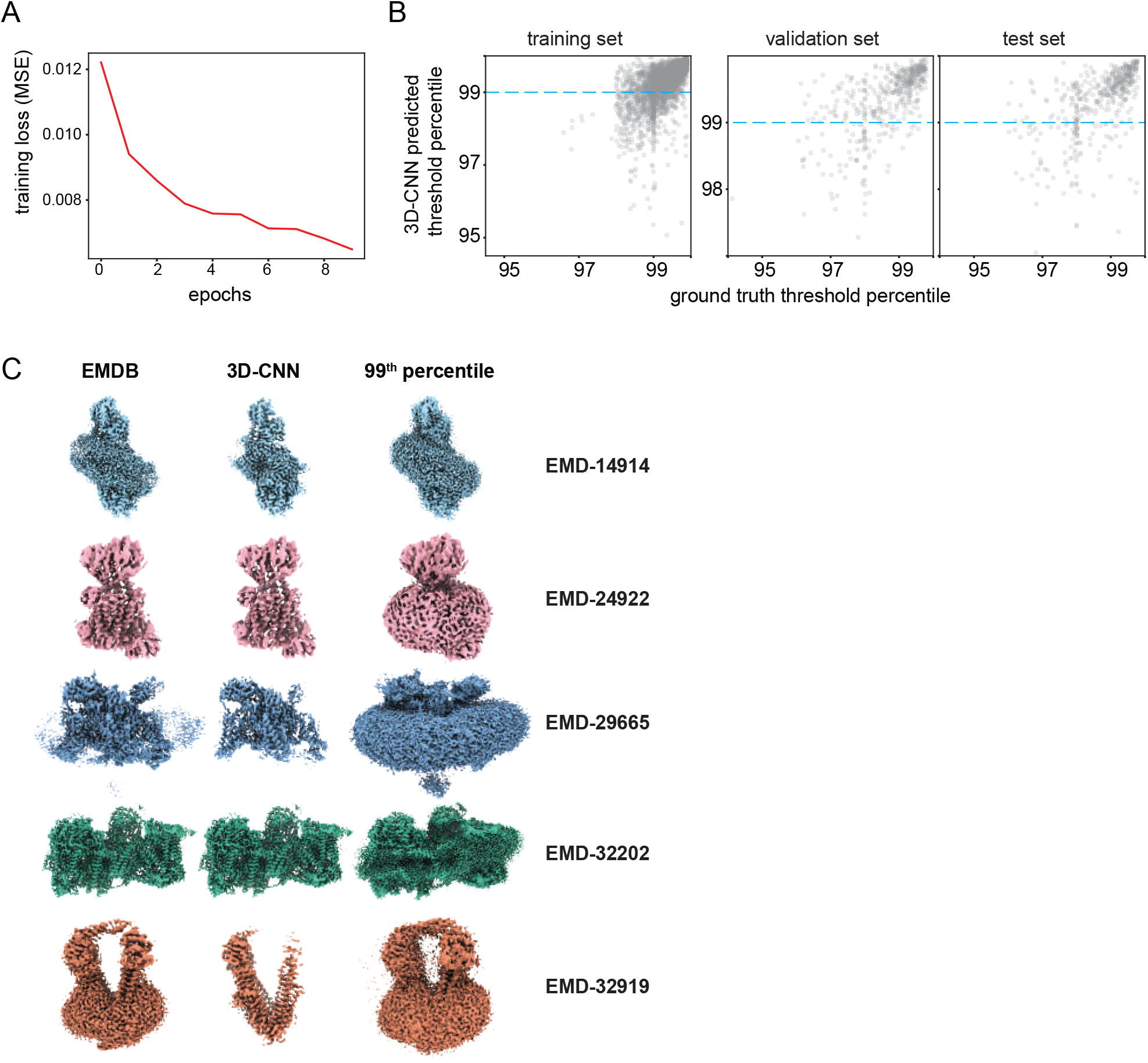
3D-CNN training, and testing. **(A)** 3D-CNN training loss (see Methods) plotted as function of number of epochs trained. **(B)** 3D-CNN predicted threshold plotted against EMDB ground truth thresholds, with each point plotted as percentile of the relevant volume data, for training, validation, and test sets. The 99th percentile of volume data, a widely-used statistical approach to predicting binarization thresholds, is shown as a dotted blue line, which highlights the significant number of volumes whose ground-truth and 3D-CNN predicted labels deviate from this value. **(C)** Representative density maps of transmembrane proteins displayed top-to-bottom from Silberberg *et al*. 2022, Shen *et al*. 2022, Huang *et al*. 2023, Gu et al. 2022, and Xiong et al. 2023, respectively. Maps are presented at binarization thresholds from EMDB ground truth (left), 3D-CNN-predicted threshold (middle), and 99th percentile threshold (right).

**Supplemental Figure 3:**
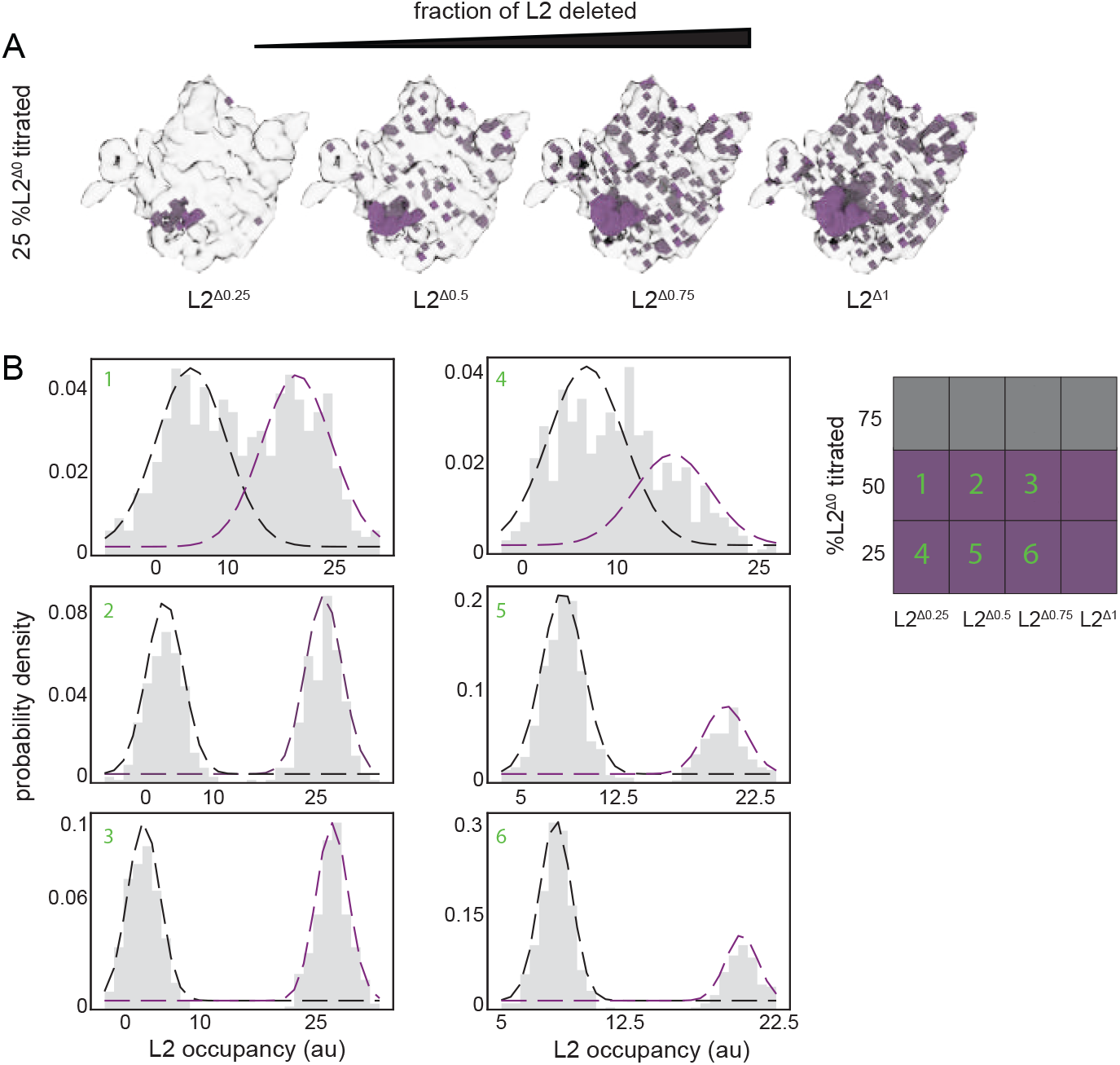
Simulated dataset of compositional heterogeneity in the *E. coli* 50S ribosome. **(A)** uL2 blocks detected by SIREn for the four datasets containing 25% uL2-intact particles. **(B)** Per-volume uL2 block occupancies for datasets containing either 50% or 25% uL2-intact particles (gray, see Methods). Datasets are labeled as shown in heatmap key. Dashed lines indicate fit Gaussian mixture model distributions (black – low occupancy class, purple – high occupancy class).

**Supplemental Figure 4:**
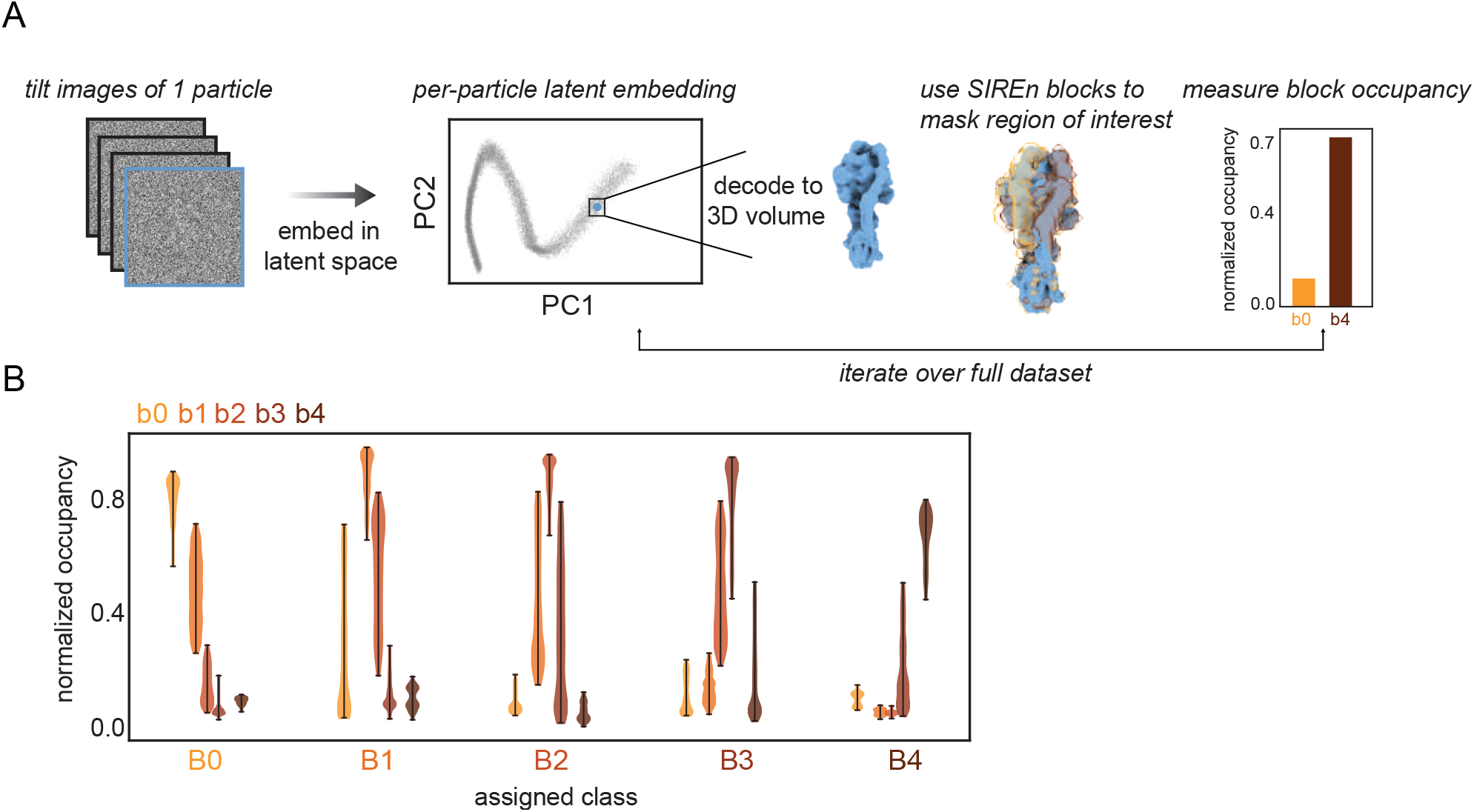
SIREn analysis of a simulated cryo-ET dataset bearing conformational heterogeneity. **(A)** Schematic workflow for ‘on-the-fly’ querying of the occupancy of SIREn blocks for each particle across a full particle stack. **(B)** Violin plots showing distributions of normalized occupancy of all five SIREn blocks in each inferred particle class (see Methods). Blocks are colored as in Figure 3.

## SUPPLEMENTARY TABLES

**Supplemental Table 1:** SIREn running times for selected datasets. Timing experiments were performed on a worker node equipped with an NVIDIA 3090 24 GB GPU and two Intel Xeon Gold 6242R CPUs (3.10GHz, 512 GB RAM), as described in the Methods. Timing experiments were performed for a variety of volume ensemble sizes and box sizes, and for datasets with a range of number of voxels within the solvent mask. All timing experiments were run using the same node and computational resources. Note that the `--filter` flag only applies to the `sketch_communities` and `expand_communities` commands.

| Dataset | Number of volumes | Box size (px) | Number of voxels in solvent mask | Command | --filter | Running time (hr:min:sec) |
| --- | --- | --- | --- | --- | --- | --- |
| ATP synthase | 500 | 64 | 7,924 | preprocess | n/a | 0:01:30 |
|  |  |  |  | eval_model |  | 0:00:04 |
|  |  |  |  | sketch_communities | no | 0:05:39 |
|  |  |  |  | expand_communities | no | 0:05:12 |
| uL2 deletion | 500 | 64 | 6,412 | preprocess | n/a | 0:00:25 |
|  |  |  |  | eval_model |  | 0:00:05 |
|  |  |  |  | sketch_communities | no | 0:03:53 |
|  |  |  |  | expand_communities | no | 0:01:57 |
| EMPIAR 10499 (intramolecular) | 500 | 32 | 6,119 | sketch_communities | no | 0:04:21 |
|  |  |  |  | expand_communities | no | 0:03:44 |
|  |  | 64 | 27,914 | preprocess | n/a | 0:00:22 |
|  |  |  |  | eval_model |  | 0:00:04 |
|  |  |  |  | sketch_communities | yes | 0:41:00 |
|  |  |  |  | expand_communities | yes | 0:25:53 |
|  |  | 96 | 75,439 | preprocess | n/a | 0:00:55 |
|  |  |  |  | eval_model |  | 0:00:04 |
|  |  |  |  | sketch_communities | yes | 3:21:41 |
|  |  |  |  | expand_communities | yes | 1:53:08 |
|  | 100 | 64 | 30,677 | preprocess | n/a | 0:00:04 |
|  |  |  |  | eval_model |  | 0:00:02 |
|  |  |  |  | sketch_communities | yes | 0:40:58 |
|  |  |  |  | expand_communities | yes | 0:00:29 |
|  | 1,000 | 64 | 27,754 | preprocess | n/a | 0:00:44 |
|  |  |  |  | eval_model |  | 0:00:07 |
|  |  |  |  | sketch_communities | yes | 0:46:42 |
|  |  |  |  | expand_communities | yes | 0:45:39 |
| EMPIAR 10499 (intermolecular) | 500 | 64 | 86,979 | preprocess | n/a | 0:00:22 |
|  |  |  |  | eval_model |  | 0:00:04 |
|  |  |  |  | sketch_communities | yes | 6:17:08 |
|  |  |  |  | expand_communities | yes | 3:43:31 |
| EMPIAR 10076 | 500 | 64 | 12,547 | preprocess | n/a | 0:00:22 |
|  |  |  |  | eval_model |  | 0:00:04 |
|  |  |  |  | sketch_communities | no | 0:10:27 |
|  |  |  |  | expand_communities | no | 0:08:03 |

**Supplemental Table 2:**
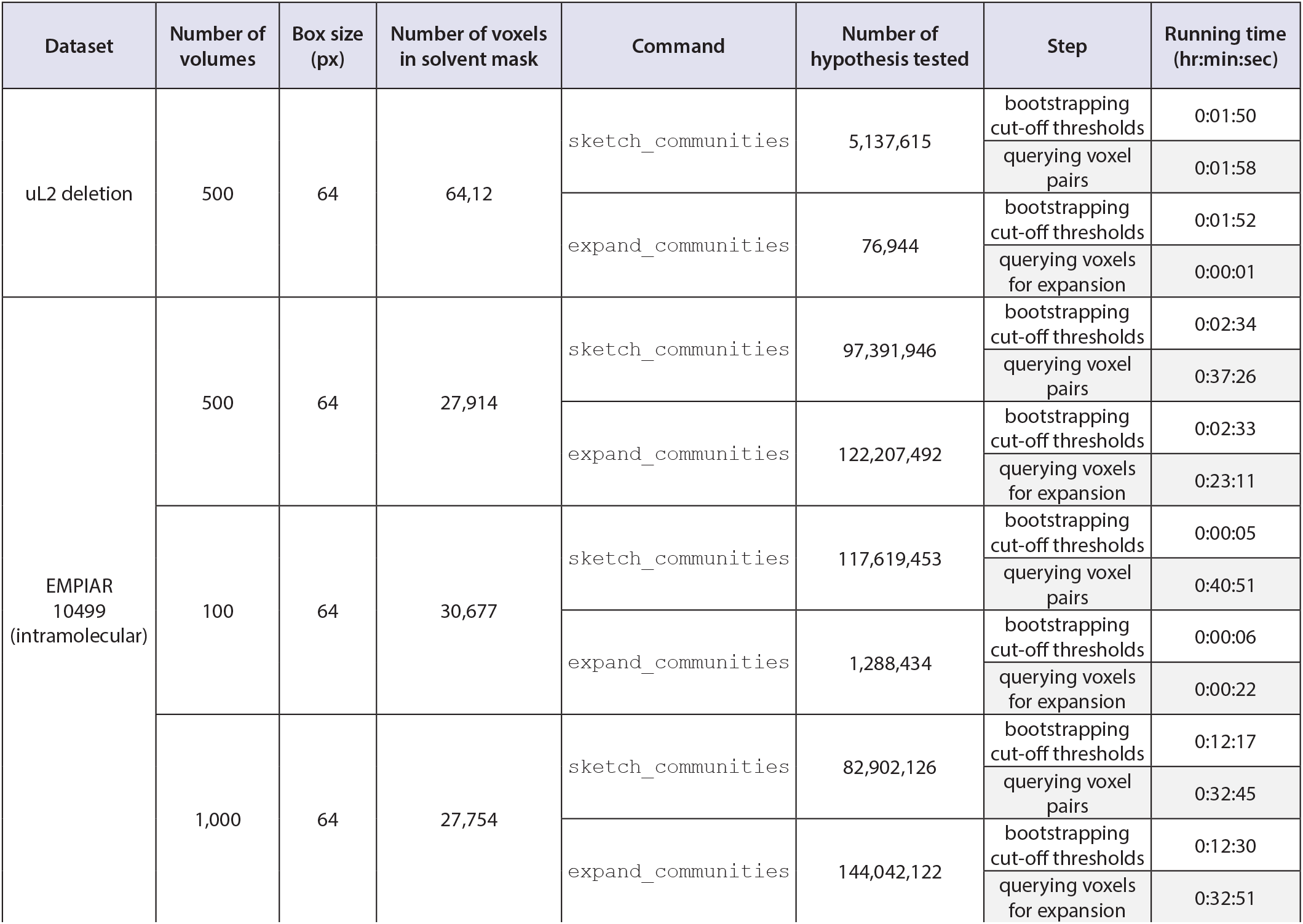
Expanded SIREn running time information. Timing information for two key steps of SIREn are presented for selected datasets. The number of hypotheses tested in the `expand_communities` command, which is calculated as the product of the number of voxels in the solvent mask, and the number of seed blocks generated by `sketch_communities`, is indicated. Note that run times depend, in part, on the number of hypotheses tested. Timing experiments were performed on a worker node equipped with an NVIDIA RTX 3090 24 GB GPU and two Intel Xeon Gold 6242R CPUs (3.10GHz, 512 GB RAM), as described in the Methods.

## REFERENCES

Baxter WT, Grassucci RA, Gao H, Frank J. 2009. Determination of signal-to-noise ratios and spectral SNRs in cryo-EM low-dose imaging of molecules. Journal of Structural Biology 166: 126–132.

Beckers M, Jakobi AJ, Sachse C. 2019. Thresholding of cryo-EM density maps by false discovery rate control. IUCrJ 6: 18–33.

Campello RJGB, Moulavi D, Zimek A, Sander J. 2015. Hierarchical Density Estimates for Data Clustering, Visualization, and Outlier Detection. ACM Transactions on Knowledge Discovery from Data (TKDD) 10.

Chen M, Ludtke SJ. 2021. Deep learning-based mixed-dimensional Gaussian mixture model for characterizing variability in cryo-EM. Nature Methods 2021 18:8 18: 930–936.

Cordasco G, Gargano L. 2010. Community detection via semi-synchronous label propagation algorithms. 2010 IEEE International Workshop on Business Applications of Social Network Analysis, BASNA 2010.

Davis JH, Tan YZ, Carragher B, Potter CS, Lyumkis D, Williamson JR. 2016. Modular Assembly of the Bacterial Large Ribosomal Subunit. Cell 167: 1610-1622.e15.

de Teresa-Trueba I, Goetz SK, Mattausch A, Stojanovska F, Zimmerli CE, Toro-Nahuelpan M, Cheng DWC, Tollervey F, Pape C, Beck M, et al. 2023. Convolutional networks for supervised mining of molecular patterns within cellular context. Nature Methods 2023 20:2 20: 284–294.

Fry MY, Najdrová V, Maggiolo AO, Saladi SM, Doležal P, Clemons WM. 2022. Structurally derived universal mechanism for the catalytic cycle of the tail-anchored targeting factor Get3. Nat Struct Mol Biol 29: 820–830.

Ghanbarpour A, Cohen SE, Fei X, Kinman LF, Bell TA, Zhang JJ, Baker TA, Davis JH, Sauer RT. 2023. A closed translocation channel in the substrate-free AAA+ ClpXP protease diminishes rogue degradation. Nature Communications 2023 14:1 14: 1–10.

Ghanbarpour A, Sauer RT, Davis JH. 2023 A proteolytic AAA+ machine poised to unfold a protein substrate. bioRxiv 2023.12.14.571662.

Ghanbarpour A, Telusma B, Powell BM, Zhang JJ, Bolstad I, Vargas C, Keller S, Baker T, Sauer RT, Davis JH. 2024. An asymmetric nautilus-like HflK/C assembly controls FtsH proteolysis of membrane proteins. bioRxiv 2024.08.09.604662.

Gilles MA, Singer A. 2024. Cryo-EM Heterogeneity Analysis using Regularized Covariance Estimation and Kernel Regression. bioRxiv 2023.10.28.564422.

Giri N, Roy RS, Cheng J. 2023. Deep learning for reconstructing protein structures from cryo-EM density maps: Recent advances and future directions. Current Opinion in Structural Biology 79: 102536.

Gu J, Liu T, Guo R, Zhang L, Yang M. 2022. The coupling mechanism of mammalian mitochondrial complex I. Nature Structural & Molecular Biology 2022 29:2 29: 172–182.

Gui M, Ma M, Sze-Tu E, Wang X, Koh F, Zhong ED, Berger B, Davis JH, Dutcher SK, Zhang R, et al. 2021. Structures of radial spokes and associated complexes important for ciliary motility. Nature structural & molecular biology 28: 29.

Guo H, Rubinstein JL. 2022. Structure of ATP synthase under strain during catalysis. Nature Communications 2022 13:1 13: 1–9.

Herreros D, Mata CP, Noddings C, Irene D, Krieger J, Agard DA, Tsai M-D, Sorzano COS, Carazo JM. 2024. Real-space heterogeneous reconstruction, refinement, and disentanglement of CryoEM conformational states with HetSIREN. bioRxiv 2024.09.16.613176.

Himes B, Grigorieff N. 2021. Cryo-TEM simulations of amorphous radiation-sensitive samples using multislice wave propagation. IUCrJ 8: 943–953.

Huang J, Fan X, Jin X, Jo S, Zhang HB, Fujita A, Bean BP, Yan N. 2023. Cannabidiol inhibits Nav channels through two distinct binding sites. Nature Communications 2023 14:1 14: 1–9.

Joseph AP, Lagerstedt I, Jakobi A, Burnley T, Patwardhan A, Topf M, Winn M. 2020. Comparing Cryo-EM Reconstructions and Validating Atomic Model Fit Using Difference Maps. Journal of Chemical Information and Modeling 60: 2552–2560.

Khavnekar S, Kelley R, Waltz F, Wietrzynski W, Zhang X, Obr M, Tagiltsev G, Beck F, Wan W, Briggs J, et al. 2023. Towards the Visual Proteomics of C. reinhardtii using High-throughput Collaborative in situ Cryo-ET. Microscopy and Microanalysis 29: 961–963.

Khavnekar S, Vrbovska V, Zaoralova M, Kelley R, Beck F, Klumpe S, Kotecha A, Plitzko J, Erdmann PS. 2022. Optimizing Cryo-FIB Lamellas for sub-5Å in situ Structural Biology. bioRxiv.

Kinman LF, Powell BM, Zhong ED, Berger B, Davis JH. 2022. Uncovering structural ensembles from single-particle cryo-EM data using cryoDRGN. Nature Protocols 2022 18:2 18: 319–339.

Lamm L, Zufferey S, Righetto RD, Wietrzynski W, Yamauchi KA, Burt A, Liu Y, Zhang H, Martinez-Sanchez A, Ziegler S, et al. 2024. MemBrain v2: an end-to-end tool for the analysis of membranes in cryo-electron tomography. bioRxiv 2024.01.05.574336.

Lee J, Lee H, Mun D. 2022. 3D convolutional neural network for machining feature recognition with gradient-based visual explanations from 3D CAD models. Scientific Reports 2022 12:1 12: 1–14.

Leesch F, Lorenzo-Orts L, Pribitzer C, Grishkovskaya I, Roehsner J, Chugunova A, Matzinger M, Roitinger E, Belačić K, Kandolf S, et al. 2023. A molecular network of conserved factors keeps ribosomes dormant in the egg. Nature 613: 712–720.

Leidig C, Thoms M, Holdermann I, Bradatsch B, Berninghausen O, Bange G, Sinning I, Hurt E, Beckmann R. 2014. 60S ribosome biogenesis requires rotation of the 5S ribonucleoprotein particle. Nat Commun 5: 3491.

LeNail A. 2019. NN-SVG: Publication-Ready Neural Network Architecture Schematics. Journal of Open Source Software 4: 747.

McInnes L, Healy J, Melville J. 2018. UMAP: Uniform Manifold Approximation and Projection for Dimension Reduction. arXiv.

Meng EC, Goddard TD, Pettersen EF, Couch GS, Pearson ZJ, Morris JH, Ferrin TE. 2023. UCSF ChimeraX: Tools for structure building and analysis. Protein Science 32: e4792.

Mu Y, Sazzed S, Alshammari M, Sun J, He J. 2021. A Tool for Segmentation of Secondary Structures in 3D Cryo-EM Density Map Components Using Deep Convolutional Neural Networks. Frontiers in Bioinformatics 1.

Nakane T, Scheres SHW. 2021. Multi-body Refinement of Cryo-EM Images in RELION. Methods in molecular biology (Clifton, NJ) 2215: 145–160.

Noble AJ. 2024. VirtualIce: Half-synthetic CryoEM Micrograph Generator. bioRxiv 2024.09.28.615520.

Noeske J, Wasserman MR, Terry DS, Altman RB, Blanchard SC, Cate JHD. 2015. High-resolution structure of the Escherichia coli ribosome. Nature Structural & Molecular Biology 2015 22:4 22: 336–341.

Pfab J, Si D. 2019. Automated threshold selection for cryo-EM density maps. ACM-BCB 2019 - Proceedings of the 10th ACM International Conference on Bioinformatics, Computational Biology and Health Informatics 161–166.

Powell BM, Brant TS, Davis JH, Mosalaganti S. 2024. Rapid structural analysis of bacterial ribosomes in situ. bioRxiv 2024.03.22.586148.

Powell BM, Davis JH. 2024. Learning structural heterogeneity from cryoelectron sub-tomograms with tomoDRGN. Nature Methods 2024 1–12.

Punjani A, Fleet DJ. 2021. 3D variability analysis: Resolving continuous flexibility and discrete heterogeneity from single particle cryo-EM. Journal of Structural Biology 213: 107702.

Punjani A, Fleet DJ. 2023. 3DFlex: determining structure and motion of flexible proteins from cryo-EM. Nature Methods 2023 20:6 20: 860–870.

Punjani A, Rubinstein JL, Fleet DJ, Brubaker MA. 2017. cryoSPARC: algorithms for rapid unsupervised cryo-EM structure determination. Nature Methods 2017 14:3 14: 290–296.

Rabuck-Gibbons JN, Lyumkis D, Williamson JR. 2022. Quantitative Mining of Compositional Heterogeneity in Cryo-EM Datasets of Ribosome Assembly Intermediates. Structure (London, England : 1993) 30: 498.

Ranno N, Si D. 2022. Neural representations of cryo-EM maps and a graph-based interpretation. BMC Bioinformatics 23: 1–20.

Rice G, Wagner T, Stabrin M, Sitsel O, Prumbaum D, Raunser S. 2023. TomoTwin: generalized 3D localization of macromolecules in cryoelectron tomograms with structural data mining. Nat Methods 20: 871–880.

Scheres SHW. 2012. RELION: Implementation of a Bayesian approach to cryo-EM structure determination. Journal of Structural Biology 180: 519.

Scheres SHW, Gao H, Valle M, Herman GT, Eggermont PPB, Frank J, Carazo JM. 2006. Disentangling conformational states of macromolecules in 3D-EM through likelihood optimization. Nature Methods 2007 4:1 4: 27–29.

Shen J, Hu M, Fan X, Ren Z, Portioli C, Yan X, Rong M, Zhou M. 2022. Extracellular domain of PepT1 interacts with TM1 to facilitate substrate transport. Structure 30: 1035-1041.e3.

Sheng K, Li N, Rabuck-Gibbons JN, Dong X, Lyumkis D, Williamson JR. 2023. Assembly landscape for the bacterial large ribosomal subunit. Nature Communications 2023 14:1 14: 1–10.

Sigworth FJ. 2016. Principles of cryo-EM single-particle image processing. Microscopy 65: 57.

Silberberg JM, Stock C, Hielkema L, Corey RA, Rheinberger J, Wunnicke D, Dubach VRA, Stansfeld PJ, Hänelt I, Paulino C. 2022. Inhibited KdpFABC transitions into an E1 off-cycle state. eLife 11.

Sitzmann V, Martel JNP, Bergman AW, Lindell DB, Wetzstein G. 2020. Implicit Neural Representations with Periodic Activation Functions. arXiv 2006.09661.

Sun J, Kinman LF, Jahagirdar D, Ortega J, Davis JH. 2023. KsgA facilitates ribosomal small subunit maturation by proofreading a key structural lesion. Nature Structural & Molecular Biology 2023 1–13.

Tan YZ, Baldwin PR, Davis JH, Williamson JR, Potter CS, Carragher B, Lyumkis D. 2017. Addressing preferred specimen orientation in single-particle cryo-EM through tilting. Nat Methods 14: 793–796.

Tang G, Peng L, Baldwin PR, Mann DS, Jiang W, Rees I, Ludtke SJ. 2007. EMAN2: An extensible image processing suite for electron microscopy. Journal of Structural Biology 157: 38–46.

Tegunov D, Xue L, Dienemann C, Cramer P, Mahamid J. 2021. Multi-particle cryo-EM refinement with M visualizes ribosome-antibiotic complex at 3.5 Å in cells. Nature Methods 2021 18:2 18: 186–193.

Turner J, Abbott S, Fonseca N, Pye R, Carrijo L, Duraisamy AK, Salih O, Wang Z, Kleywegt GJ, Morris KL, et al. 2024. EMDB—the Electron Microscopy Data Bank. Nucleic Acids Research 52: D456–D465.

Wang J, Li D, Chen L, Cao W, Kong L, Zhang W, Croll T, Deng Z, Liang J, Wang Z. 2022. Catalytic trajectory of a dimeric nonribosomal peptide synthetase subunit with an inserted epimerase domain. Nat Commun 13: 592.

Wang JY, Tuck OT, Skopintsev P, Soczek KM, Li G, Al-Shayeb B, Zhou J, Doudna JA. 2023. Genome expansion by a CRISPR trimmer-integrase. Nature 618: 855–861.

Xiong C, Jia LN, Xiong WX, Wu XT, Xiong LL, Wang TH, Zhou D, Hong Z, Liu Z, Tang L. 2023. Structural insights into substrate recognition and translocation of human peroxisomal ABC transporter ALDP. Signal Transduction and Targeted Therapy 2022 8:1 8: 1–13.

Xue L, Lenz S, Zimmermann-Kogadeeva M, Tegunov D, Cramer P, Bork P, Rappsilber J, Mahamid J. 2022. Visualizing translation dynamics at atomic detail inside a bacterial cell. Nature 2022 610:7930 610: 205–211.

Zhang C, Cantara W, Jeon Y, Musier-Forsyth K, Grigorieff N, Lyumkis D. 2019. Analysis of discrete local variability and structural covariance in macromolecular assemblies using Cryo-EM and focused classification. Ultramicroscopy 203: 170–180.

Zhong ED, Bepler T, Berger B, Davis JH. 2021. CryoDRGN: reconstruction of heterogeneous cryo-EM structures using neural networks. Nature Methods 18.

Zivanov J, Nakane T, Forsberg BO, Kimanius D, Hagen WJH, Lindahl E, Scheres SHW. 2018. New tools for automated high-resolution cryo-EM structure determination in RELION-3. eLife.

